# Response of sleep slow oscillations to acoustic stimulation is evidenced by distinctive synchronization processes

**DOI:** 10.1101/2022.05.28.493849

**Authors:** Miguel Navarrete, Alejandro Osorio-Forero, Angela Gómez, David Henao, Michel Le Van Quyen, Mario Valderrama

**Affiliations:** Department of Biomedical Engineering, University of Los Andes, Cra. 1 N° 18A–12, Bogotá, Colombia; Department of Fundamental Neurosciences, University of Lausanne, Rue du Bugnon 9, CH-1005 Lausanne, Switzerland; Department of Psychology, University of Los Andes, Cra. 1 N° 18A–12, Bogota, Colombia; Laboratoire d’Imagerie Biomédicale (LIB), U1146 INSERM- SU - CNRS 7371, Campus des Cordeliers, 15 rue de l’Ecole de Médecine, Paris, France

**Keywords:** slow waves, slow oscillations, K-complex, sleep, auditory stimulation, closed loop stimulation, NREM, synchronization

## Abstract

Closed loop acoustic stimulation (CLAS) during sleep has shown to boost slow wave (SW) amplitude and spindle power. Moreover, sleep SW are suggested to be classified based on different processes of neuronal synchronization. Following this, different types of SW events may have distinct functional roles and be differentially affected by external stimuli. However, the SW synchronization processes affected by CLAS are not well understood. Here, we studied the effect of CLAS on the dissociation of SW events based on two features of neuronal synchronization in the EEG (topological spread and wave slope). We evaluated and classified individual SW events of fourteen healthy subjects during a CLAS stimulated (STM) and a control night (CNT). Three main categories of SW events were found denoting (C1) steep-slope SW with global spread, (C2) flat-slope waves with localized spread and homeostatic regulation, and (C3) multipeaked flat-slope events with global spread. Comparing between conditions, we found a consistent increase of event proportion and trough amplitudes for C1 events during the time of stimulation. Furthermore, we found similar increases in post-stimulus spectral power in θ, β and σ frequencies for CNT vs STIM condition independently of sleep stage or SW categories. However, topological analysis showed differentiated spatial dynamics in N2 and N3 for SW categories and the co-occurrence with spindle events. Our findings reveal the nature of post-stimulus SW and suggest that CLAS boosts SW amplitudes by increasing neuronal synchronization of wave troughs, leading thus the post-stimulus SW-spindle co-occurrence.

## INTRODUCTION

Slow wave sleep (SWS) is characterized by large amplitude cortical slow wave activity (0.5-4 Hz) in the electroencephalogram (EEG). Large slow waves (SW) are generated by neocortical neurons which slowly alternate between phases of membrane depolarization associated with increased firing (up-states) and hyperpolarization associated with reduced firing (down-states) (Achermann and Borbély, 1997; Steriade et al., 1993a). Furthermore, the corticothalamic interactions that occur during SW give rise to faster oscillatory events: thalamic sleep spindles (SS: 0.5-3s amplitude events in the sigma band 11-16 Hz) (Axmacher et al., 2008; Le Van Quyen et al., 2016, 2010). The coupled interaction between these activities and hippocampal sharp-wave ripples (130–250 Hz, SWR) is thought to play a key role for memory consolidation (Inostroza and Born, 2013; Klinzing et al., 2019; Rasch and Born, 2013).

Although the nature of SW as an active process is still debated, not all dynamics during SW events may contribute equally to the cognitive function they are often associated with (Crunelli et al., 2018). Therefore, a dichotomy of SW has been proposed in the literature dissociating these events by their frequency range (Steriade et al., 1993c), sleep pressure (Hubbard et al., 2020), cholinergic (Nghiem et al., 2020) and thalamic drive (Fernandez et al., 2018), roles in memory consolidation (Kim et al., 2019) and recruitment of neuronal population (Bouchard et al., 2021). Following these, recent findings suggest that the function of SW could be defined their synchronization characteristics. Hence, EEG synchronization during SW synthesize the similarity of spatial-temporal fluctuations of neuronal firing in both local and distant cortical regions, thus driving both the characteristic frequency and the spatial spreading of SW (Kim et al., 2019; Siclari et al., 2014). Specifically, synchronization of local circuits has been related to the SW slope and the lowest-to-highest SW amplitude transition (Bouchard et al., 2021; Vyazovskiy et al., 2009), whereas spatial spreading of SW may indicate the modulation of neuronal bi-stability (firing and silence) across brain regions (Nir et al., 2011). Hence, faster delta oscillations (1 – 4Hz) originate from the cortico-cortical networks and the thalamus, while the larger and wider slow oscillations (SO: 0.5 – 2 Hz) are thought to arise from cortico-cortical activations but strengthened by thalamic dynamics (Lemieux et al., 2014; Steriade et al., 1993b). Both delta waves and SOs can nest spindles, possibly by increasing the occurrence of cortical down-states on the thalamus during SW troughs (Mak-McCully et al., 2017). However, these different waves may also have distinct neuronal functions. Specifically, intra-thalamic GABAergic synapses are potentiated or depressed depending on whether the corticothalamic neurons are modulated by SOs or delta rhythms (Crunelli et al., 2018; Kim et al., 2019). Likewise, the global dynamics of SW (i.e., whether they are localized or widespread) seem determine their involvement in physiological (Nir et al., 2011) and learning processes (Fattinger et al., 2017) as well as they can respond differently to sleep pressure (Hubbard et al., 2020). Consistently, SW were suggested to be classified by their synchronization efficiency – synchronization strength determined by the wave slope and spread –, which dissociates two distinct processes (Bernardi et al., 2018; Siclari et al., 2014). From these studies, SW were characterized as large, widespread, and steep “Type I” waves, or smaller, flatter and more localized “Type II” waves. Specifically, these ongoing SW synchronization dynamics are thought to reflect specific cortico-cortical and corticothalamic neuronal dynamics, which may have different functional roles and to be differentially affected by external stimuli (Bernardi et al., 2018).

Consistent with the notion that the brain is more excitable during specific slow wave phases (Bergmann et al., 2012; Schabus et al., 2012), some studies evidenced that auditory stimuli applied around the peak of the SW provokes a strong intensification of SW amplitude and associated spindle activity (Ngo et al., 2013; Ong et al., 2016). This closed-loop acoustic stimulation (CLAS) has also been reported to boost declarative memory performance (Ngo et al., 2013; Ong et al., 2016; Papalambros et al., 2017). In this sense, boosting SW is perhaps beneficial because it increases the coupling between cortical SOs, corticothalamic spindles and high-frequency hippocampal SWR during SWS (Maingret et al., 2016; Staresina et al., 2015). However, the sole increase of post-stimulus SW amplitude might not be enough to boost memory consolidation (Henin et al., 2019). Indeed, the cognitive improvements induced by CLAS are known to be limited by age (Schneider et al., 2020) and the type of memory test implemented (Leminen et al., 2017). Presumably, CLAS might only enhance SO-SS-SWR coupling when the stimulus is able to increase the corticothalamic coordination of SW troughs which then sustain the co-occurrence of cortical down-states in the thalamus (Navarrete et al., 2020b; Todorova and Zugaro, 2020). Indeed, the capability of enhancing this corticothalamic coordination may not only lie in the effect of the stimulation, but also in the ongoing dynamics of the cortical activity during the CLAS click (Navarrete et al., 2022, 2020a). However, currently it is not clear how the stimulus disturbs the ongoing neuronal dynamics and the characteristics of the resulting post-stimulus SW.

Consequently, in this work we focused on characterizing the underlying SW types from their EEG synchronization attributes across the night: spread and main slope. Additionally, we evaluated how CLAS modulates these synchronization features in the post-stimulus SW. We hypothesized that CLAS modulates SWS by controlling the ongoing synchronization dynamics of the post stimulus SW events, which then leads to changes of wave co-occurrence with associated spindles. Specifically, we first developed and validated a heuristic approach to classify SW depending on their synchronization attributes. Then, we evaluated the response to in-phase acoustic stimulation and measured these features during the manipulation. We chose an adaptive approach to deliver real-time sound stimuli targeting peak-phases of SOs. This approach took into consideration the relatively long time of slow waves which allowed us to examine a wide range of electrophysiological responses in terms of their frequency and topography. We found that CLAS facilitated larger, synchronous, and global cortico-cortical SOs, possibly sustained and reinforced by thalamic dynamics. Interestingly, spindles events were more associated with post-stimulus SW with strengthened EEG synchronization. We demonstrate that CLAS modulates SWS by enhancing the cortical SW synchronization which primarily leads to larger SOs and cooccurring post-stimulus spindle events.

## MATERIALS AND METHODS

### Scalp EEG subjects and recordings

Fourteen healthy, free of medication, and non-smoker volunteers (mean age 26 ± 4 years, nine women) were recruited at the University of Los Andes in Bogotá, Colombia for the study. All subjects were asked to avoid caffeine, energetic and alcoholic drinks 24 h before the experiment and not to take naps on the day of the study. Participants gave their written informed consent and procedures were approved by the ethical committee of the University of Los Andes. Each subject underwent two recording nights, one for stimulation (STM) and one for control (CNT, using muted sounds). The order of experimental conditions was randomized across subjects and separated by at least one week. Night sessions consisted of polysomnographic recordings performed in line with the recommendations from the American Academy of Sleep Medicine (AASM) (Iber et al., 2007). The electrodes included: 2 electrooculogram (EOG) channels (right and left eyes), 2 electromyogram (EMG) channels of the submentalis muscle, and 21 EEG channels in the standard international 10-20 system (Fp1, Fpz, Fp2, F7, F3, Fz, F4, F8, T3, C3, Cz, C4, T4, T5, P3, Pz, P4, T6, O1, Oz, O2) which were all referenced to the linked mastoids (A1 and A2). Data were recorded with a LTM64 Micromed System (Treviso, Italy) and signals were sampled at 256 Hz.

### Sleep and EEG analysis

Sleep was visually scored by a reviewer blinded to the stimulation condition and according to the AASM criteria (Iber et al., 2007). All hypnograms were visually confirmed by a time frequency analysis (Figure S1).

Slow wave activity (SWA) was obtained from the EEG signals filtered between 0.5 - 2 Hz using a zero-phase windowed equiripple FIR filter (3 dB at 0.25 and 3.08 Hz; <-37 dB at f < 0.01 Hz and f > 4 Hz). SW were considered when their negative deflection had consecutive zero crossings between 0.25 s and 1.4 s, regardless of the wave amplitude (Riedner et al., 2007). This procedure was used to facilitate the detection of both local and global SW (Mensen et al., 2016).

Spindle activity was computed by applying a zero-phase bandpass FIR filter between 11 – 16 Hz (3 dB at 10.62 and 17.38 Hz, <-40 dB at f < 10.01 Hz and f > 18 Hz). Then, the root mean squared (RMS) was calculated using a time window of 0.2 s (Clemens et al., 2007). Candidate spindle events were detected as the RMS activity larger than a threshold established as the 86.64 percentile (equivalent to 1.5 SD for a Gaussian distribution) of the spindle activity during NREM and with durations between 0.3 and 3 s (Warby et al., 2014). Furthermore, events were required to have at least five cycles, a unimodal peak in the spindle frequency band (11 – 16 Hz) and decreasing power for higher frequencies computed by the Morlet wavelet (Purcell et al., 2017). Stimuli-dependent sleep spindles were defined as the events beginning in the trough-to-trough interval next to the marked click (Navarrete et al., 2020a).

NREM sleep cycles were determined as the beginning of any sustained N2 or N3 period (undisturbed >30s) until the beginning of either REM or awakening. To account for NREM cycle variability between nights, we evaluated the first three NREM cycles independently for each subject, but fourth and any further NREM cycles were collapsed into one group. Therefore, NREM cycle analyses include four groups (1st NREM cycle, 2nd NREM cycle, 3th NREM cycle, and 4th+ NREM cycle). Event density was evaluated as the number of events by minute, then changes in SW density for each category were z-scored by subject and compared within the same NREM cycle.

### Offline slow wave classification

For the classification of SW, we implemented a heuristic method based on two slow wave synchronization characteristics: topological spread and wave slope (Riedner et al., 2007; Siclari et al., 2014). We computed the synchronization characteristics of each slow wave based on measures proposed in a previous study using high-density EEG (Bernardi et al., 2018), but slightly modified to take into account the lower resolution of our recordings and the variability of more localized slow wave events. As described in Figure 1a, we first set a unique time reference across channels by computing a virtual channel defined as the grand average of the SWA over all EEG electrodes. Then, we identified SW events in this virtual channel which were further used as reference events (rSW). Briefly, rSW were identified as negative deflections with consecutive zero crossings between 0.25 to 1.0 seconds. Additionally, we further restricted the amplitude of the averaged SWA to less than −5 μV to minimize the effect of random oscillations across channels. Next, for each channel, we identified SW for which the negative half-wave was within 100 ms from the rSW event. Hence, we computed the mean slope (*λ*) of all detected SW across channels. Similarly, we assessed the trough synchronization by computing the regularized SW scalp involvement (*ε*). For computing *ε*, we first built a distribution of the number of channels with overlapping negative half-waves detected across time indicating the wave involvement (W_i_). Then, *ε* was calculated as the normalized area under the curve of the W_i_ distribution (Figure 1a Top).

**Figure 1.**
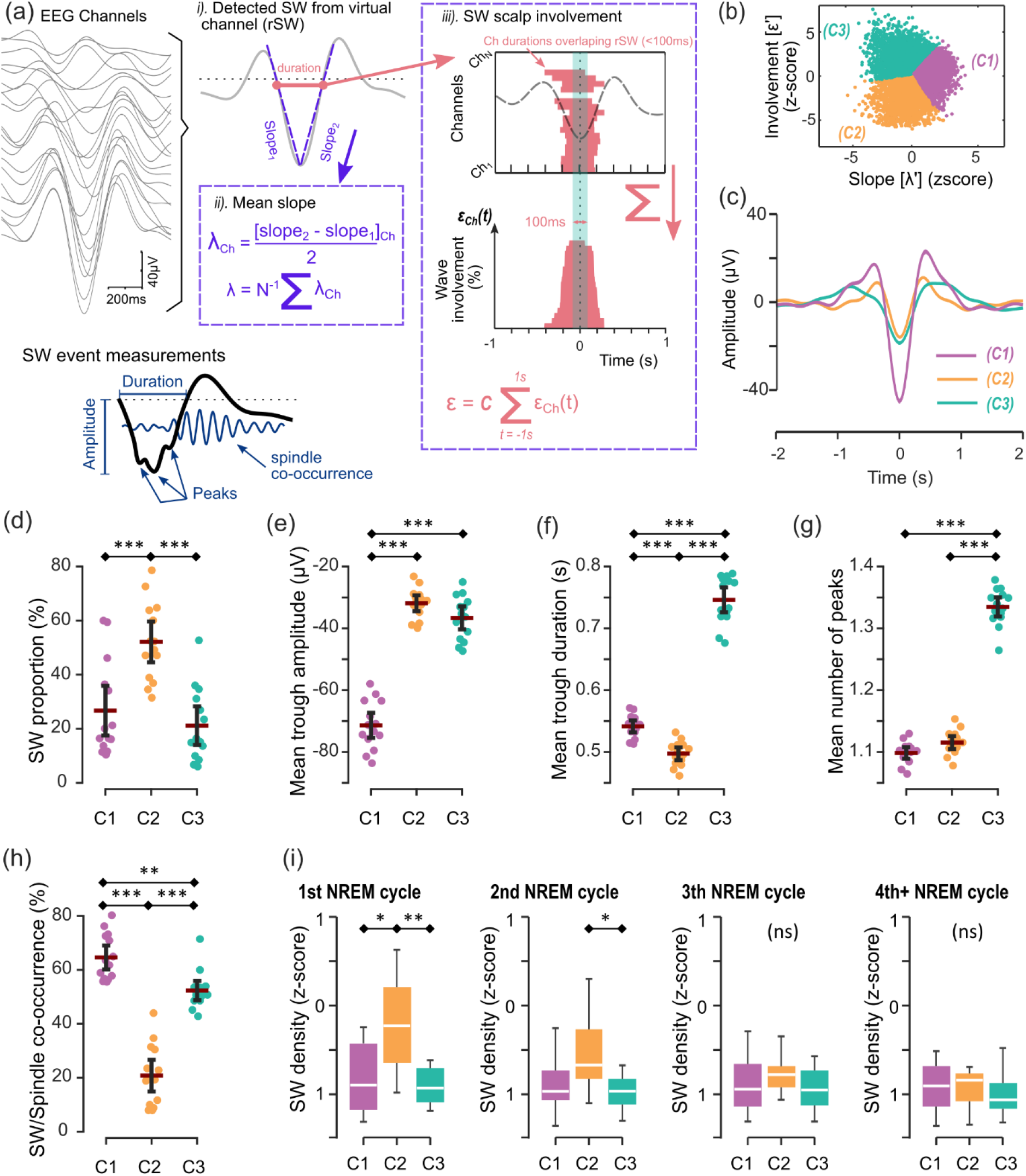
Classification and characteristics of spontaneous SW categories. a) Features of slow wave synchronization were computed for characterizing different types of SW. For each detected slow wave, we determined the average slope across scalp channels (λ). Furthermore, we computed the wave involvement as the area under the curve from the normalized time distribution of overlapping negative half-wave durations across scalp channels (ε). b) Regularized and z scored λ and ε values (λ’ and ε’) were then used as inputs for an unsupervised k-means classifier. Three different clusters were thus determined from this data featuring three different SW categories. c) The trace average for the resulting SW categories sorted over the largest trough: C1 indicating large and steep SOs, C2 showing small waves, and C3 indicating long durations SW troughs. Differences in characteristics between spontaneous SW categories averaged across electrodes in (d) event proportion, (e) trough amplitude, (f) trough duration. (g) number of negative peaks, (h) co-occurrence with detected sleep spindles, and (i) homeostatic decline.

Subsequently, we classified SW depending on their *λ* and *ε* characteristics. In this sense, previous studies suggest the use of a unidimensional *synchronization score* to classify SW (Bernardi et al., 2018; Siclari et al., 2014). Nevertheless, a unidimensional *synchronization score* can overshadow slow wave dynamics between these two categories. For this reason, we chose to keep the bidimensionality in our classification process. Thus, we used a k-means algorithm to group SW events into three categories as suggested by a Silhouette analysis computing the optimal number of classes (Figure S3a). For this, first, we log-transformed the mean slope (*λ*) and applied a square-root transformation to the regularized scalp involvement (*ε*) to homogenize the data variance. We then normalized these values using a z-score transformation before the k-means classification (Milligan and Cooper, 1988) (Figure S3b). The k-means algorithm was trained using all detected spontaneous SW across all night from the CNT condition in all subjects. Then, using the same z-score parameters from spontaneous events, SW from the detection/stimulation CLAS periods were normalized and assigned to the corresponding categories by finding the k-nearest neighbor from the classified spontaneous waves.

We computed multiple morphological measures characteristic of the identified SW categories. These measures include the event proportion, trough amplitudes, trough duration, number of peaks, and SW/spindle co-occurrence. The event proportion was determined as the percentage of SW events of each category from all detected SW. Trough measures were determined as the average amplitudes and time durations of the negative half wave across channels for each identified SW event. Furthermore, we also identified the average number of peaks – across channels – for each detected SW trough in the SWS frequency band (0.5 – 4 Hz). Finally, we determine the SW/spindle co-occurrence as the percentage of SW troughs associated with a subsequent spindle (Figure 1a Bottom).

### Auditory stimulation

Acoustic stimuli consisted of stereophonic pink noise clicks with a 50 ms duration and a rising and falling slopes lasting 5 ms. Precise timestamps were recorded online when the clicks were applied. The sound stimulus was applied using AcousticSheep SleepPhones^®^ headphones specially designed for sleep, and the volume was individually set for each participant to a non-disturbing level. For each night, the stimulation protocol started when the subject achieved sustained N2 or N3 sleep stages and stopped on any sign of arousal. Stimulation was applied during the two first NREM cycles only. Following previous protocols (Ngo et al., 2015, 2013; Ong et al., 2016), two consecutive auditory clicks were targeted to the peaks of ongoing SOs through a closed-loop stimulation system. For this, an algorithm for real-time automatic detection and stimulation was developed (Figure S2a). Briefly, the EEG signal from a reference electrode (F3) was pre-processed online by applying a low-pass Chebyshev window-based FIR filter with cut-off of 9 Hz at 3 dB and < −40 dB for frequencies in the stopband above 17 Hz. Then, all negative deflections (i.e., under a negative amplitude threshold of −60 μV) were evaluated for further online processing. If both the time of the detected negative deflection (t_zz_: from positive-to-negative to negative-to-positive zero crossings) and the peak-to-trough time (t_PN_) were within a time interval corresponding to the duration of a half-wave of the SO frequency band (~250 – 1400 ms), then the detected deflection was considered an ongoing SO trough (Figure 2a, Figure S2a). Next, detected ongoing SOs were either used as reference-waves or as stimulation-waves (Figure S2a). Specifically, reference-waves were not stimulated but used to compute the click-time (*t_s_*). For this, the peak-time (*t_p_*) was determined as the timelapse between the negative-to-positive zero crossing and the subsequent SO peak, and this was saved to memory. The *t_p_* times were classified into fifteen different half-wave t_zz_ time-range intervals equally spaced logarithmically between 250 and 1400 ms. Then, the click-time *t_s_* was computed as the average of the last three *t_p_* times from the previous reference-waves within the same t_ZZ_ time-range of the detected SO trough (Figure S2b). Hence, for simulation-waves, the click was applied at *t_s_* measured from the negative-to-positive zero crossing following the detected SO trough. Following previous recommendations for maximal performance (Ngo et al., 2015), a second click was applied on the subsequent wave with *t_s_’* computed as before (Figure 1b, Figure S2a). Then, a refractory time of 2.5 seconds without any stimulation was determined after the last click. All SOs detected during this refractory non-stimulated period were thus used as reference-waves.

**Figure 2.**
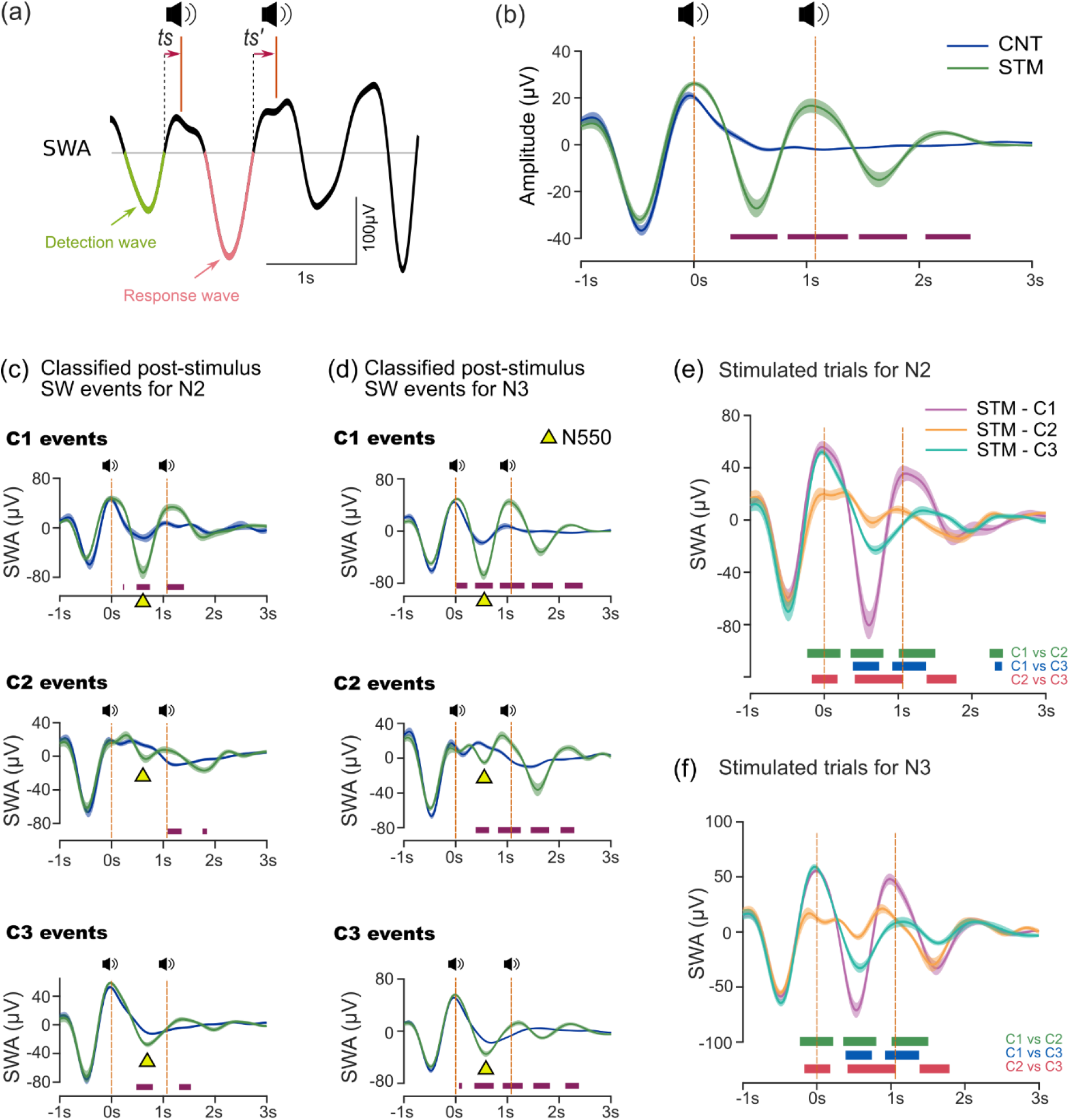
CLAS stimulation pos-stimulus SW categories. a) large SW were online detected based on amplitude and wave duration features. The time for stimulation (*t_s_*) was applied from the negative-to-positive zero-crossing and computed as the average of time-to-peak from previous unstimulated SW. Two consecutive clicks were applied, and post-stimulation negative half waves were used for further analysis. b) The typical response to CLAS suggest the enhancement of post-stimulus SO amplitudes when comparing STM with CNT conditions. Average response to stimulation after organizing post-stimulus slow-wave categories for c) N2 and d) N3. An increment of amplitude average was evident for all post-stimulus wave types indicating an ERP N550 component (yellow triangle). The axes illustrate ERP differences between CNT and STM conditions for post-stimulus C1 events (top), C2 events (top), and C3 events (bottom). A comparison of average trials for the different post-stimulus slow wave categories are displayed for e) N2 and f) N3. These ERP responses show the characteristics for the different SW categories.

Experimental protocols for STM and CNT conditions were similar, but click volumes were muted in the CNT condition. All algorithms were implemented using custom-made MATLAB GUI (Mathworks, Natick, MA, USA), and the Psychophysics Toolbox extensions for the acoustic stimulation (Brainard, 1997; Kleiner et al., 2007; Pelli, 1997).

For trial analyses, only stimulations during N2 and N3 sleep stages were evaluated. Stimuli that were within less than 2 s from any arousal or artifact were removed. The amplitude response of the SWA to the auditory click was estimated as the absolute trough amplitude following the acoustic click. Consequently, all trials in which the subsequent stimuli were wrongly placed before the subsequent SO trough were excluded from the analysis.

### Statistical Analyses

Significant differences between STM vs CNT conditions for pairwise comparisons across subjects were obtained using two-tailed paired-samples t-tests. Difference scores (Δ) between conditions were computed by subtracting CNT measures from STM. Thus, a positive score indicated a preference for STM condition, and a negative for the CNT condition. Multiple testing problem of unidimensional measures was corrected using the Bonferroni correction. One-way analyses of variance (ANOVA) were implemented to compare morphological measures between categories. Tukey’s honestly significant difference procedure was implemented as a multiple comparison post-hoc test. A repeated measures ANOVA was performed to compare the effect of NREM cycles across the night on the density of SW categories.

For time and frequency analyses, between-subject trial statistics for STM vs CNT conditions were performed using nonparametric cluster-level statistics (Bullmore et al., 1999; Maris and Oostenveld, 2007). Suprathreshold clusters were determined by adjacent Welch’s t-values which absolute value was larger than the 98^th^ percentile from the permutation distribution of the maximal statistic (Nichols and Holmes, 2002). Cluster-level statistics were obtained from the distribution of the maximal suprathreshold cluster size after 1600 non-repeated permutations (Butar et al., 2008; Keller-Mcnulty and Higgins, 1987). A Benjamini-Yekutieli procedure for controlling the false discovery rate (FDR) was applied to correct for multiple comparisons in multi-channel analyses (Benjamini and Yekutieli, 2001).

Finally, data was expressed as mean ± 95% confidence intervals (CI) for comparisons across subjects. Statistics were considered significant at *p* < 0.05 after correcting for multiple comparisons. All statistical analyses were performed in MATLAB.

## RESULTS

### Sleep architecture

We evaluated the sleep macro-structure between conditions of stimulation (Control vs Stimulation). Consistently with previous studies using CLAS, Table S1 shows that we did not find significant differences in the sleep macro-structure between these two conditions (Leminen et al., 2017; Ngo et al., 2013; Schneider et al., 2020). These results suggest that CLAS do not interfere in the sleep continuity and the stimulation did not induce a larger time of awakenings.

### Classification of slow waves for spontaneous SWS

We first analyzed the differences of SW events based on their synchronization features. First, we classified spontaneous SW into different categories depending on their synchronization characteristics, i.e. mean slope (*λ*) and regularized SW scalp involvement (*ε*) (Figure 1a). These features were normalized and regularized before the classification process (Figure S3b). A Silhouette analysis indicated that the best number of categories is 3 (silhouette value of 0.585) (Figure S3a). Although the slope vs involvement plot did not indicate obvious grouping of the events, we discerned a three-cornered distribution of the data which followed the characteristics of the resulting categories (Figure 1b). SW were thus classified in three categories: Category 1 (C1) corresponding to events with the largest slope and high involvement, Category 2 (C2) events with the lowest involvement and small to middle slope values, Category 3 (C3) events with largest involvement and low slope (Figure 1c).

To better understand the characteristics of the identified categories, we evaluated the morphological wave characteristics of the identified groups in spontaneous SWS. One-way ANOVA analyses indicated main effects in event proportion (F = 16.39, p < .001, Figure 1d), trough amplitude (F = 146.24, p < .001, Figure 1e), trough duration (F = 339.63, p < .001, Figure 1f), mean number of peaks across all channels (F = 460.93, p < .001, Figure 1g), and spindle co-occurrence (F = 87.97, p < .001, Figure 1h). Specifically, post-hoc analyses showed an increased proportion of all-night C2 events when compared to C1 and C3 waves (both p < 0.001, Figure 1d). Next, we found that the average trough amplitudes were much larger for C1 events compared to both C2 and C3 (both p < 0.001 Figure 1e). Similarly, we found that the mean trough duration was considerably larger for C3 when compared to C1 and C2 groups, as well as C1 trough duration was also larger than C2 (all p < 0.001, Figure 1f). The mean number of peaks of C3 was larger than both C1 and C2 (both p < 0.001, Figure 1g). Finally, for spindle co-occurrence, we found significant differences across the three categories (Figure 1h), with a larger proportion of SS events co-occurring with the total number of C1 events when compared to C2 and C3 (both p < .001) as well as a larger proportion of SS events co-occurring with C3 compared to C2 waves (p = .002). Similar results were evidenced when analyzing N2 (Figure S4) and N3 (Figure S5) sleep stages independently.

We also evaluated the probabilities of SW transition between categories for sequences of events (events < 2s spaced out). The three SW categories represent nine different possible transitions in which one subsequent wave could be the same or different category. An one way ANOVA showed that there are differences between the probabilities of transitions (F(7, 118) = 32.66, p < .001). A post-hoc analysis indicated the largest probability for a sequence of events represented by C2-to-C2 transitions (~43.73%) which had significant differences compared with all other combinations, whereas the lower probability was for C1-to-C3 transitions (~3.58%) (Figure S6).

Next, we evaluated the differences of SW density by each NREM cycle and across the night (Figure 1i). A repeated measures ANOVA indicated that there was a statistically significant effect of time on total SW density (F(3, 90) = 6.38, p < .001), and a significant effect of time on SW density by category (F(6, 90) = 2.66, p = .020). Then, when evaluating differences within each sleep cycle, one-way ANOVA analyses indicated main effects in SW density in the 1^st^ NREM cycle (F = 12.05, p < .001) and the 2^nd^ NREM cycle (F = 3.78, p = .030), but not for the 3^th^ NREM cycle (F = 2.51, p = .095) or the 4^th^+ NREM cycle (F = 0.44, p = .645). In the 1^st^ NREM cycle, post hoc analyses indicated larger C2 event density when compared with both C1 (p = .002) and C3 (p < .001) event densities. Instead, C2 event density was only larger when compared with C3 event density (p = .025) for the 2^nd^ NREM cycle.

In this sense, here we identified categories of spontaneous SW that gracefully fit sleep graphoelements described in previous works. Thus, previous studies have characterized SW as large, widespread, and steep “type I” waves or smaller, flatter and more localized “type II” waves in humans (Bernardi et al., 2018; Siclari et al., 2014) or the delta 1 and delta 2 waves described in rodents (Hubbard et al., 2020). In keeping with this, the identified categories indicated (C1) steep slope SW with global spread (Type I – like waves), (C2) localized flat slope SW (Type II – like waves). Furthermore, we found a third category (C3) typifying flat slope SW with global spread (sharing characteristics of both type I and type II waves). Hence, C1 and C3 slow wave events signpost stereotypical characteristics of SOs (Iber et al., 2007), C2 events are consistent with the morphology of local SW (Riedner et al., 2007), with a further homeostatic decline characteristic of delta activity (Hubbard et al., 2020). However, C3 events may group heterogeneous activity from multiple cortical sources including both large global SOs as well as more localized activities (Chavez et al., 2006; Hubbard et al., 2020).

### Slow wave categories in response to CLAS

Next, we aimed to investigate the SW in response to CLAS. For this, we implemented a protocol of stimulation considering the time variability of SO peaks (Figure 2a). Our algorithm of SO detection and stimulation was accurately stimulating at a mean phase of −0.15 ± 0.58 radians (Figure S2c) where 0 radians denotes SO peak and ±π/2 radians are the zero crossings. Furthermore, our algorithm had an online SO detection accuracy of 84.06 ± 8.42% (ratio of offline detected and stimulated SOs vs total clicks). Figure 2b shows how the stimuli increased the after-click SW amplitude, resulting in a succession of consecutive SW which were larger than those following the non-stimulated trials. Thus, our protocol of stimulation showed a cortical response to CLAS which enhances SW as described in the literature. (Ngo et al., 2013; Ong et al., 2016).

We compared the morphological features of SW categories across all the night and between conditions (Table S2). We did not find differences for morphological features of each SW category between CNT or STM conditions when comparing all the events across the night (all p > 0.057). Similarly, we did not find differences for event density for each NREM cycle between CNT and STM nights in none of the three SW categories (all p > 0.08).

Next, we evaluated the different characteristics of the evoked SW after the stimulus click. Independently of the slow wave category, Event related potential (ERP) analyses showed an increase in trough amplitude for STM vs CTN condition in both N2 and N3 stages (Figure 2c for N2 and Figure 2d for N3). Specifically, the ERP analysis averaged across all slow wave categories showed an induced N550 component as the main post-stimulus trough (~578 ms ± 55 ms after the click). This increased trough shows thus the characteristic SO enhancement previously shown in CLAS works (Ngo et al., 2013; Ong et al., 2016; Schneider et al., 2020). However, we also found differences in the post-stimulus SWA when comparing the STM response between categories in both N2 and N3. Hence, C1 events presented larger amplitudes in response to CLAS, C3 negative deflections had longer durations, and C2 events resulted in shorter and flatter wave averages (all p < 0.05, after FDR correction, Figure 2e-f).

Then, we evaluated the post-stimulus synchronization caused by CLAS. A two-way ANOVA was performed to analyze the effect of post-stimulus SW category (3 types) and sleep stage (2 types) on the difference score Δ(%) of the proportion of post-stimulus events (Figure 3a). We found a significant interaction between the effects of the sleep stage and SW category (F(2, 78) = 4.55, p = .013). Simple main effects analysis showed that the sleep stage did not influence the difference score of the percentage of events (p = 1), but the event category did show an effect (p < .001). For both N2 and N3, post-hoc analyses showed an increase in the percentage of events for C1 caused by CLAS, whereas the percentage of C2 and C3 events decreased (all C1 vs C2 and C1 vs C3, p < .001).

**Figure 3.**
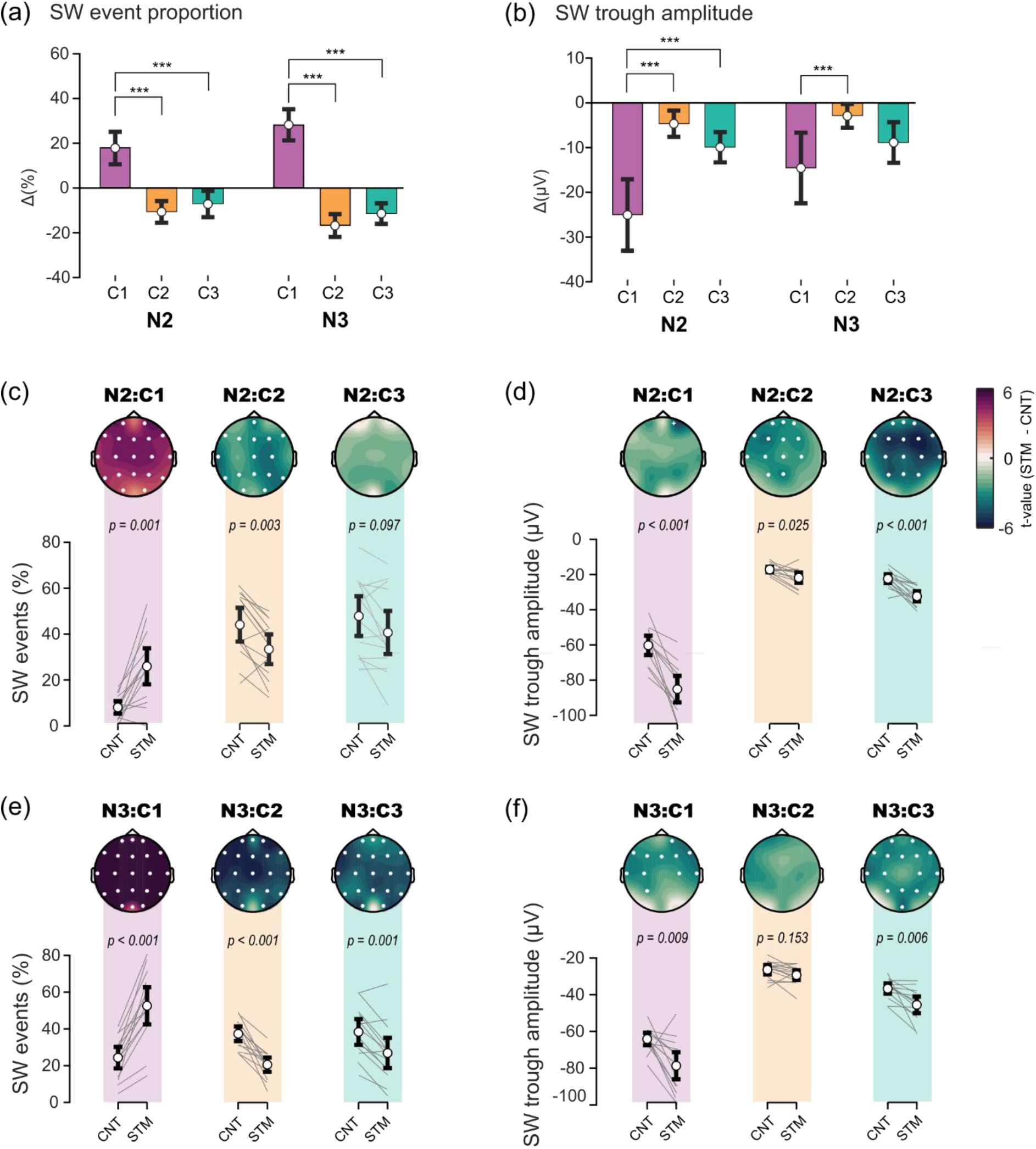
Changes in SW categories induced by CLAS. a) The difference score (Δ) of the percentage of SW events in each category indicated an increased ratio of C1 events while the proportion of C2 and C3 events decreased in both N2 and N3. b) larger difference score increments of negative trough amplitudes were evident for C1 and C3 events in both sleep stages. c) Changes in SO proportion were global for all SW categories in N3. d) Significant increments of slow wave amplitude were evident for C1 events in frontocentral channels and in whole brain channels for C3 events in N3. e) Changes in SO proportion were also global for C1 and C2 SW categories in N2. f) Significant increments of SW amplitude included frontal, central and parietal channels in N2 for C2 and C3 events. White dots in topoplots indicate channels with significant differences after FDR correction.

Similarly, for CLAS responses we analyzed the effect of post-stimulus slow wave category and sleep stage on difference score Δ(μV) of SW trough amplitudes (Figure 3b). The two-way ANOVA analysis revealed no interaction between the sleep stage and category (F(2, 75) = 1.06, p = .351). Likewise, simple main effects analysis showed no effect of sleep stage (p = .081), but the category did have an effect on the difference score for event troughs (p < .001). Specifically, a post-hoc analysis showed larger C1 troughs compared to respective C2 and C3 troughs (both p < .001) for stimulated trials during N2. However, for stimuli applied in N3, only C1 troughs had larger increments when compared to C2 (p < .001) but not when compared to C3 (p = .127).

Then, we evaluated the topographic characteristics for the between-condition changes in slow wave categories. Consistent with the previous analyses, for clicks in N2 we found a stimulation-induced increase in the number of C1 events and a decrease of C2 in all channels. C3 events for N2 showed a decrease of event proportion but these differences did not survive after FDR correction (Figure 3c). However, we found a stimulation-induced increase in the number of C1 events as well as a decrease of both C2 and C3 events during N3 in all channels (Figure 3e).

Next, we evaluated topographic differences in SW trough amplitudes across stimulation conditions. For N2, we found significant changes in SW troughs between stimulation conditions in all categories. Although most differences between channels did not survive FDR correction for C1 events, an enhancement of SW trough amplitudes was evident in frontal, central and parietal channels in resulting C2 and C3 events (Figure 3d). Instead, for N3, no differences in trough amplitude were found for C2 events. However, we found a frontocentral increase of trough amplitudes for C1 events and a global increase of trough amplitudes for C3 events (Figure 3f).

Thus, our results indicate that CLAS induces an increase of N550 components across all resulting SW categories and sleep stages. Nevertheless, stimuli did induce changes in the synchronization dynamics (SW slope and spread) for resulting SW by increasing C1 events in both N2 and N3, but with different topographic responses for C2 and C3 events between sleep stages. Additionally, event amplitudes are larger for frontal C1 while the increase of C3 event amplitudes are more globally spread when comparing CNT and STM trials in N3.

### Spectral response to CLAS between slow wave categories

We also investigated the spectral response to CLAS for each SW category. First, we evaluated the time-frequency response to CLAS between 1 – 30 Hz for all channels and trials. This analysis showed increased spindle activity, as described in previous works (Ngo et al., 2015; Ong et al., 2016) (Figure 4a). We found three main time-frequency clusters of increased spectral power when evaluating the differences between CNT and STM conditions. These frequency clusters highlighted an increase in the theta band (θ cluster: 3 – 9.6 Hz, ~0 – 875 ms), sigma (σ cluster: 11.4 – 15.6 Hz, ~687 – 1406 ms) and beta (β cluster: 16.2 – 23.2 Hz, ~312 – 797 ms) frequencies after the stimulus (Figure 4b, top left).

**Figure 4.**
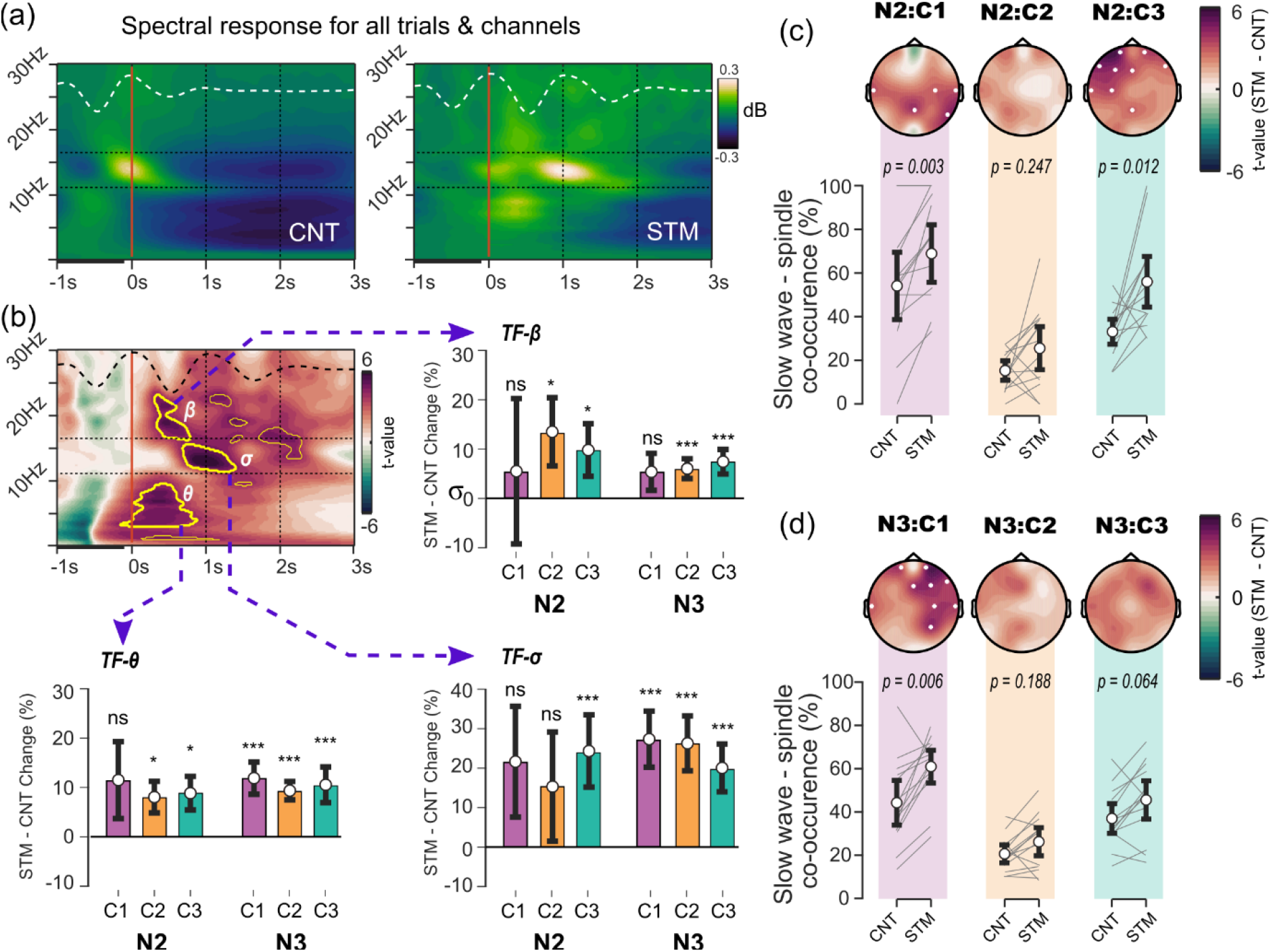
Post-stimulation spectral changes depending on resulting slow wave categories after CLAS. (a) Average spectral response across channels and subjects in CNT and STM conditions. (b) Cluster differences of spectral response between CNT and STM (significative clusters in yellow border). Three large spectral clusters were identified by cluster-based statistics. No significant differences of changes in spectral power were found between sleep stages or between the resulting post-stimulus slow wave categories. (c) CNT-to-STM changes in spindle co-occurrence indicated an increased proportion on frontocentral channels contralateral to the stimulus for resulting C1 events during N3. Changes in proportion of spindle co-occurrence was not significant for C2 and C3 events (d). Instead, during N2 the spindle co-occurrence increased for post-stimulus parietal C1 events and frontal C3 events. White dots in topoplots indicate channels with significant differences after FDR correction.

Next, we evaluated the spectral changes in these frequency clusters of interest (θ, σ and β) for different slow wave categories after CLAS. First, we evaluated the difference scores for cluster power in each condition. We found a consistent increase θ power in response to CLAS during N3 for all slow wave categories (C1, p < .001; C2, p < .001, C3, p < .001), but only C2 and C3 categories survived Bonferroni correction in N2 (C1, p = .113; C2, p = .002, C3, p = .002) (Figure 4b, bottom left). Similarly, a significant increase of σ power after CLAS was found for all slow wave clusters in N3 (C1, p < .001; C2, p < .001, C3, p < .001), but this increase was only significant for C3 in N2 (C1, p = .087; C2, p = .439, C3, p = .001) (Figure 4b, bottom right). Instead, for β power we observed a significant increase after CLAS for C2 and C3 slow wave clusters in both N3 (C1, p = .131; C2, p < .001, C3, p < .001) and N2 (C1, p > .999; C2, p = .019, C3, p = .029) (Figure 4b, top right). Secondly, we applied two-way ANOVAs to evaluate the difference score in spectral power depending on sleep stage and slow wave categories. However, we did not find effects of either sleep stage or SW category for the cluster-frequencies of interest. All together, these results suggest that CLAS induces changes in relative power of θ, σ and β, although no power differences were found neither across sleep stages nor between post-stimulus SW categories.

### Spindle co-occurrence between slow wave categories during CLAS

Finally, we detected sleep spindles and analyzed the SW-spindle co-occurrence after CLAS for each slow wave category. During N2 (Figure 4c), we found that spindle co-occurrences with C1 events increased in parietal regions (p = .003). Similarly, differences between conditions were found for spindle co-occurrence with C3 events (p = .012), and these differences were mostly found in frontal electrodes. Nevertheless, we report no significant findings when considering spindle co-occurrence with C2 events (p = .247). Conversely, during N3 (Figure 4d), we found increased SW-spindle co-occurrence in the right frontocentral channels for STM vs CNT condition for C1 events (p = .006). No differences were found in SW-spindle co-occurrence for neither C2 nor C3 slow wave categories (C2, p = .188; C3, p = .064).

In the previous section, the power cluster analysis showed an increase of σ power regardless of the SW categories, but this result was not the same for detected spindles. Instead, detected spindle-events in response to CLAS were mostly facilitated after evoked C1 waves across sleep states, and for C3 events during N2.

## DISCUSSION

Here we report a systematic investigation of the slow wave dynamics and distribution of brain responses to CLAS during NREM sleep. Our data is in line with previous studies, confirming that CLAS can effectively enhance slow wave amplitudes, together with theta and sigma activity (Leminen et al., 2017; Ngo et al., 2013; Ong et al., 2016; Papalambros et al., 2017; Schneider et al., 2020). Using an unsupervised method, we classified SW into three groups depending on their synchronization characteristics (scalp spread and wave slope). Thus, SW were grouped as: (i) C1: large, widespread, and steep SW, (ii), C2: small and localized waves with homeostatic decline, and (iii) C3: widespread, small, flat slope and multipeaked SW. We found that CLAS induced more synchronized and widespread SWs (C1) but decreased the likelihood of more localized or multi-formed events (C2, C3). Nevertheless, all post-stimulation waves showed increased ERP amplitudes and increased θ, σ and β power, regardless of the evaluated sleep stages.

### Slow wave categories during unstimulated NREM sleep

Our wave classification method successfully discriminated morphological synchronization characteristics consistent with SW types previously described. The patterns categorized here are similar to SW types that characterize the early and late epochs of the falling asleep period (Siclari et al., 2014) and in the stable NREM sleep (Bernardi et al., 2018). Specifically, C1 events here described fit the stereotypical description of SOs and K-complexes (Iber et al., 2007). Similar to previously described δ1-activity in rodents, C1 waves lack of homeostatic decay, lie in the frequency range of low delta (~0.5 – 2.75 Hz) and convey a global contribution of multiple cortical regions (Hubbard et al., 2020). Similarly, in line with the previously described Type I SW in humans (Siclari et al., 2014), C1 events are also characterized by their low density, large amplitude, steep slope and long duration. As Type I events, C1 events could be mediated by the arousal-promoting structures. Indeed, highly synchronized thalamus-driven cortico-cortical NREM oscillations enforce synchronized SOs, differentiating these from the non-thalamic-driven SW in terms of function, extension, and neuronal synchronization (Crunelli and Hughes, 2010). Nevertheless, C1 events did not present microarousals at the scalp level which are typical of K-complexes, thus suggesting more stable dynamics of their neuronal synchronization mechanisms. C2 waves entailed smaller and more localized events with similar characteristics to those previously identified as regional SW (Nir et al., 2011). Similarly, these represent events in faster frequency ranges with homeostatic decline as well as high delta (δ2-activity: ~2.75 – 4 Hz) (Hubbard et al., 2020). Spontaneous C2 events showed a larger proportion of SW with smaller amplitudes, flat slope and short duration, such as Type II SW (Siclari et al., 2014). Also, C2 slow wave events were less synchronous across scalp electrodes and showed lower co-occurrence with SS. Hence, the more untangled SW-spindles dynamics for C2 events suggest lower corticothalamic interactions during these events (Mak-McCully et al., 2017; Navarrete et al., 2020b) although their homeostatic decline suggest a possible thalamic involvement in their origin as for δ2-activity (Hubbard et al., 2020). Additionally, we identified a third category grouping SW events that could be regarded as a middle point between both C1 and C2 types. This third group here denoted as the C3 events included widespread, small, flat slope and multipeaked SW. C3 events are thus likely to describe SW that share characteristics of both Type I and Type II oscillations and include much slower oscillations. Because of their wide spreading, multipeaked negative deflection, and spindle-locking characteristics, C3 SW may include nested C1 and C2 events from multiple cortical sources (Chavez et al., 2006) as well as δ1 nests δ2-activity (Hubbard et al., 2020).

Because of their characteristic spread, slope, and homeostatic decline, we claim that C1 events may correspond to the previously described δ1/Type I SW while the C2 events may correspond to δ2/Type II SW. Nevertheless, we did not use the same labels for our wave categorization because of the differences in the classification procedure as well as in the associated higher frequencies (either α or σ) co-occurring with the SW. Specifically, we found that C1 events are tightly linked with spindles. Their close co-occurrence was observed in both N2 and N3 sleep stages. This observation is in keeping with corticothalamic observations indicating that the convergence of multiple cortical large down-states on the thalamus leads the corticothalamic spindle dynamics and hence the SW-spindle co-occurrence (Mak-McCully et al., 2017). Conversely, Type I events are more readily associated with cortical arousals (particularly α activity) while Type II waves were previously linked to the relative power increase within the σ frequency (Bernardi et al., 2018). We suggest that these main differences between SW characteristics are caused by the process of wave classification. Specifically, strictly considering only two classes of slow wave could have overlooked the oscillatory events that share characteristics with other types of events. Indeed, we found a third class of SW events (C3) that might be considered as an intermediary SW type or the temporal overlap of the two first ones. This oscillatory activity was highly related to spindles, but these could have been previously considered as Type II waves. Certainly, some wave characteristics might suggest similarities between C3 events and Type II waves. For instance, they both have slightly higher event proportions, lower amplitudes, and flatter slopes than C1 waves. Likewise, multipeaked SOs such as C3 events have been suggested to predominate during late sleep periods (Riedner et al., 2007), which is consistent with the characteristics of type II waves (Bernardi et al., 2018; Genzel et al., 2014). In this sense, previous work describing SW by their synchronization characteristics may have grouped C2 and most of C3 events as Type II SW.

### Changes on slow wave categories induced by CLAS

During CLAS, the proportion of C1 events increased, suggesting a rise of cortical synchronization sustained in a stronger corticothalamic loop generated by the sound click. Stimulus-promoted C1 events are large in amplitude and highly synchronous which suggests an interaction between cortical and thalamic neurons (Bellesi et al., 2014; Navarrete et al., 2020b). Specifically, typical SOs originate from cortico-cortical activations, but they are fully expressed in amplitude, extent, and synchrony only with active thalamic contribution (Crunelli et al., 2018). Therefore, the increase of stereotypical SO events during CLAS has been suggested to be promoted by sensory-generated corticothalamic K-complexes (Krugliakova et al., 2020; Ngo et al., 2013). Indeed, K-complexes are singular events which originate from the same corticothalamic circuitry that drives the neuronal dynamics of SOs (Amzica and Steriade, 1997; Steriade, 2003). We do not rule out that post-stimulus C1 waves might be boosted by a mechanism analogous to K-complexes, but overlapping and different to the spontaneous sleep oscillations. However, our findings suggest that post-stimulus waves are not merely K-complexes that were time sorted with the stimulation. Compared to spontaneous, evoked K-complexes do not increase in amplitude (Bastien and Campbell, 1994; Campbell, 2010). Therefore, if CLAS responses were only formed by evoked K-complexes, no changes in SW amplitude would be evident, neither for the stereotypical SOs nor for the more localized waves.

Akin to the large evoked SOs, we found that C1 events increased in proportion in all channels, while the increments in trough amplitude for these SW were found mostly across the frontocentral channels. These patterns follow the topographic dynamics of both stereotypical SO and K-complexes (Bastien and Campbell, 1994; Nir et al., 2011). This is coherent with the fact that hyper synchronization seems to be facilitated by the phase entrainment and the additive responses evoked by the stimulus (Sauseng et al., 2007). Furthermore, contrary to C1 waves, the likelihood of C2 and C3 events decreased. Hence, more localized SW may have been reinforced by a hyper-synchronization driven by the impact of the stimulus on the corticothalamic networks (Lemieux et al., 2014; Rosanova and Timofeev, 2005). Also, the decrease in C3 events might represent a decrease in the number of overlapping SW from different cortical sources. However, trough amplitudes of C3 events were also enhanced by the stimulus, suggesting an additive effect caused by the click. Thus, the post-stimulus C3 events might include sensory-generated K-complexes which failed to globally synchronize the endogenous local or widespread SW.

Our findings also indicate that effects of the sound click are not absolute and may be limited by the endogenous SO activity. A previous hypothesis offered by Bellesi et al. (2014) suggests that the characteristics of the SOs evoked by CLAS depend on the conditions of the arousal system and the thalamic relay cells at the time of the stimulation. Specifically, this hypothesis explains the SO enhancement by thalamic mediated arousal (SO-TMA). This SO-TMA hypothesis proposes that SOs may be enhanced by the stimulus when thalamic matrix cells activate the associative cortex and in time windows when the noradrenergic (NA) neurons are suppressed (Bellesi et al., 2014). This hypothesis further suggests the existence of some non-explicit cortical conditions that determine cortical sensitivity to the stimulation (i.e., whether the stimulation would boost SOs dynamics, whether it would be completely ineffective, or even induce awakenings). In line with this hypothesis, recent analyses suggest that the amplitude of SO peaks may facilitate global synchronization of post-neuronal down-states, thereby enhancing the amplitudes of the resulting slow wave troughs (Navarrete et al., 2022; Torres et al., 2021; Wei et al., 2020). Congruently, we found that large and synchronous C1 and C3 events were increased by stimuli during large peaks of the SOs (with average amplitudes larger than +40 μV across all EEG channels, Figures 2e & 2f). Instead, the after-click response of C2 events showed smaller trough amplitudes when stimuli were applied during low amplitude SO peaks. Hence, the phase entrainment of the spontaneous SWA stimulated at large amplitude SO peaks drive ERP dynamics that might create the required conditions to promote C1 events.

### Effects of CLAS on the arousal response in induced slow wave categories

We found similar ERP and spectral responses to the stimulus regardless of the induced slow wave category or sleep stage. Firstly, a marked N550 ERP component was evident following CLAS. Previous studies suggest that this component may play a role as a protective mechanism against large arousals (Campbell, 2010; Halász, 2016). Furthermore, the N550 amplitude has been described to be unaffected by the intensity of the stimulus and was only noticeable for trials eliciting K-complexes (Bastien and Campbell, 1994). Nevertheless, our findings suggest that N550 could be an endogenous response of the arousal system even in the absence of the stereotypical K-complexes. Specifically, we also noticed this N550 component in post-stimulus C2 events that do not represent neither conventional SOs nor K-complexes. Therefore, this N550 component could be regarded as a phase entrainment caused by CLAS, which facilitates the large post-stimulus response and that also indicates an activation of the arousal mechanism during induced SW. Secondly, we found that the stimulation induces a power increase of β, θ, and σ when compared to the control condition. Previous works described the temporal association of these rhythms and report a localized frontal θ – β coupling that is enhanced by CLAS (Krugliakova et al., 2020). We extend the existing literature by also showing that the power induced by the stimulation did not differ neither between sleep stages nor between slow wave categories.

In keeping with the SO-TMA hypothesis, the induced ERP and spectral activity (β, θ, and σ power) may be part of the endogenous sleep protection response, coordinated by the arousal system. Indeed, the spectral response to CLAS that we report here may be mediated by the decreased activation of NA neurons during the timing of the stimulus. Specifically, the cortico-coerulear interactions that regulate cortical excitability are minimal during SOs peaks (Eschenko et al., 2012), these coinciding with the target of the acoustic clicks during CLAS. The reduced NA activity prevents the awakenings, thereby allowing the arousal system to prevent any disruptions to SWS continuity. In this sense, our results could indicate activation of the endogenous arousal system to prevent awakenings and facilitating the cortical activation of stereotypical SO or K-complex events. In this sense, the SW response to CLAS might be highly dependent on the NA tone which is evidenced by the infraslow oscillatory pattern of σ activity (Hayat et al., 2020; Lecci et al., 2017; Osorio-Forero et al., 2021). Future studies might evaluate the relationship of NA levels and acoustic stimulation by coupling the auditory stimulus to the infraslow fluctuation in σ activity.

### Co-occurrence of click induced spindles and slow wave categories

Our findings suggest that post-stimulus SW-spindle co-occurrence could depend on the corticothalamic drive of each induced SW category. Sleep spindles are oscillatory events that originate within the thalamic networks during NREM (Contreras and Steriade, 1996; Steriade, 2003). Previous work suggests that spindles generation is facilitated by the synchronous coordination between cortical down-states and thalamic neurons (Jiang et al., 2019a, 2019b; Mak-McCully et al., 2017). Specifically, concurrent cortical down-states during SW drive thalamic down-states (Crunelli and Hughes, 2010), which then hyperpolarize thalamic cells and thus trigger the thalamic spindles. These thalamic spindles are then projected to the cortex coinciding with the up-states of the ongoing SOs (Lüthi, 2014; Mak-McCully et al., 2017). Consequently, the high synchronization of cortical down-states induced by the stimulus may sustain stronger spindle activity co-occurring with post-stimulus slow wave peaks. However, the enhancement of spectral activity in the spindle band does not provide sufficient evidence for the existence of spindle events (Fernandez and Lüthi, 2020). Specifically, not all the post-stimulation SW categories showed an increment in spindle events although the σ power was boosted across sleep stages and SW events. The surge in σ power without an increase in detectable spindles might indicate an activation of thalamic arousal projected throughout the diffused thalamic matrix system (Hagler et al., 2018), or more local spindles going on in the LFP signals that cannot reach the scalp (Lacourse et al., 2019). In this sense, only the C1 events during N3 and C1 and C3 events during N2 showed both a σ power increase and an increase in co-occurring SW-spindle events after the CLAS click. This may further suggest distinctive dynamics of each sleep stage. Specifically, sleep spindles are mostly driven by stereotypical trains of SOs during N3. Here, C1 events represent SW with the largest hyper-synchronization of cortical down-states, consistent with the stereotypical SOs which are strongly coupled with sleep spindles (Klinzing et al., 2016). Thus, C1 waves may be associated with moments of reticular thalamic bursting activity related to increased SO and sigma (Fernandez et al., 2018) compared to moments when these neurons drift away from this activity pattern. Conversely, K-complexes which show similar characteristics as the C3 events, are the hallmark of N2 and the main drivers of sleep spindles during this stage. It has also been suggested that K-complexes may originate from the cortical disruption of the thalamic spindling networks (Mak-McCully et al., 2014). Hence, the distributed activation of sensory-evoked K-complexes may overlap with the ongoing local down-states, thus producing the characteristic C3 events and their coupling with spindles. On the other hand, we found a decreased likelihood of C2 and spindle events co-occurrence, and this was not significantly enhanced by the auditory click. This is consistent with previous literature demonstrating the exclusive dynamics of spindle and delta waves (Steriade et al., 1993b). Briefly, sleep spindles and delta waves could be originated within the same thalamic circuitry. Hence, scalp incidence of these events might include spindle or delta activity rather than both together (Fernandez et al., 2018).

### Study limitations

In this study we evaluated different types of SW and how the phase-locked acoustic stimulation drives SWA. Nevertheless, some precautions should be taken when interpreting our results. Firstly, the number of different wave types was determined a priori and only based on the statistical distribution of the detected spontaneous SW. Visual inspection of the data did not show a clear separation between wave categories. In this sense, wave events close to the thresholds between different clusters may share very similar characteristics with each other, given no visually evident thresholds between categories. Hence, these events may share characteristics common to one or two classes and slightly change the category upon acquisition of new observations. Nonetheless, group level characteristics are in keep with previous descriptions of the differences between SW types, even when small changes in intra-cluster thresholds are provoked by new data (Bernardi et al., 2018; Nir et al., 2011; Siclari et al., 2014). Secondly, we did not evaluate events according to their origin, but rather using a computational method based on the spatial synchronization characteristics of SW visible on the scalp EEG. In this sense, we were unable to investigate the neuronal function and neuronal dynamics involved in the generation of the resulting categories. For this, only the simultaneous recordings of cortical and sub-cortical activity could truly discern between cortical and thalamic origins. However, the observed differences between categorized slow wave events are consistent with the hypothesized mechanisms of cortico-cortical and cortico-thalamic dynamics during SWS (Crunelli et al., 2018). Thirdly, our findings did not consider interindividual variability when neither characterizing the slow wave categories nor their responses to the stimulus. Our analyses imply that the variability of slow wave categories is consistent between subjects, and this is in keep with previous works indicating global dynamics during SWS (Lemieux et al., 2014; Mukovski et al., 2007; Ngo et al., 2015; Nir et al., 2011).

### Conclusions

We differentiated diverse categories of SW during NREM based on their synchronization characteristics, wave slope and spatial spread, at scalp EEG level. We also evaluated the CLAS-induced slow waves and described the synchronization characteristics of the cortical response to the acoustic stimuli. Hence, we showed that the features of post-stimulus SW may suggest an increase of cortical activity characteristic of stronger corticothalamic synchronization during NREM sleep. Furthermore, we showed that induced higher frequencies are independent of the type of slow wave responses, suggesting an activation of the arousal system. Instead, post-stimulus spindle events were facilitated by the large, widespread, and steep SOs were evoked by the stimulation. This observation can be interpreted under the light of the consequences of CLAS in more widespread dynamics, at the cost of diminishing local interactions. Our findings help to elucidate the nature of the cortical response to the auditory stimulus during NREM sleep. Hence, future works may help to provide a better understanding of the role of the different SW types on memory consolidation, sleep recovery, sleep quality and the possible use of CLAS as a treatment for improving sleep health.

## Acknowledgments

We gratefully acknowledge funding from COLCIENCIAS call for grants 567 and MINCIENCIAS call for grants 848. The authors would like to thank Martyna Rakowska and Anne Koopman for their helpful comments and discussion on the manuscript. Likewise, we would like to thank Daniela Gonzalez and Brayan Arias for their help in the completion of the experiments. Finally, we thank all the participants for their time and commitment to the study.

## Notes

### Competing Interest Statement

The authors have declared no competing interest.

## REFERENCES

Achermann P, Borbély AA (1997) Low-frequency (< 1 Hz) oscillations in the human sleep electroencephalogram. Neuroscience 81:213–22.

Amzica F, Steriade M (1997) Cellular substrates and laminar profile of sleep K-complex. Neuroscience 82:671–686.

Axmacher N, Haupt S, Fernández G, Elger CE, Fell J (2008) The role of sleep in declarative memory consolidation - Direct evidence by intracranial EEG. Cereb Cortex 18:500–507.

Bastien C, Campbell K (1994) Effects of rate of tone-pip stimulation on the evoked K-Complex. J Sleep Res 3:65–72.

Bellesi M, Riedner B a., Garcia-Molina GN, Cirelli C, Tononi G (2014) Enhancement of sleep slow waves: underlying mechanisms and practical consequences. Front Syst Neurosci 8:1–17.

Benjamini Y, Yekutieli D (2001) The control of the false discovery rate in multiple testing under dependency. Ann Stat 29:1165–1188.

Bergmann TO, Mölle M, Diedrichs J, Born J, Siebner HR (2012) NeuroImage Sleep spindle-related reactivation of category-specific cortical regions after learning face-scene associations. Neuroimage 59:2733–2742.

Bernardi G, Siclari F, Handjaras G, Riedner BA, Tononi G (2018) Local and Widespread Slow Waves in Stable NREM Sleep: Evidence for Distinct Regulation Mechanisms. Front Hum Neurosci 12:1–13.

Bouchard M, Lina JM, Gaudreault PO, Lafreniret A, Duba J, Gosselin N, Carrier J (2021) Sleeping at the switch. Elife 10:1–17.

Brainard DH (1997) The Psychophysics Toolbox. Spat Vis 10:433–436.

Bullmore ET, Suckling J, Overmeyer S, Rabe-Hesketh S, Taylor E, Brammer MJ (1999) Global, voxel, and cluster tests, by theory and permutation, for a difference between two groups of structural MR images of the brain. IEEE Trans Med Imaging 18:32–42.

Butar FB, Ph D, Park J (2008) Permutation Tests for Comparing Two Populations. J Math Sci Math Educ 3:19–30.

Campbell K (2010) Event-related potentials as a measure of sleep disturbance: A tutorial review. Noise Heal 12:137.

Chavez M, Besserve M, Adam C, Martinerie J (2006) Towards a proper estimation of phase synchronization from time series. J Neurosci Methods 154:149–160.

Clemens Z, Molle M, Eross L, Barsi P, Halasz P, Born J (2007) Temporal coupling of parahippocampal ripples, sleep spindles and slow oscillations in humans. Brain 130:2868–2878.

Contreras D, Steriade M (1996) Spindle oscillation in cats: the role of corticothalamic feedback in a thalamically generated rhythm. J Physiol 490:159–79.

Crunelli V, Hughes SW (2010) The slow (less than1 Hz) rhythm of non-REM sleep: a dialogue between three cardinal oscillators. Nat Neurosci 13:9–17.

Crunelli V, Larincz ML, Connelly WM, David F, Hughes SW, Lambert RC, Leresche N, Errington AC (2018) Dual function of thalamic low-vigilance state oscillations: Rhythm-regulation and plasticity. Nat Rev Neurosci 19:107–118.

Eschenko O, Magri C, Panzeri S, Sara SJ (2012) Noradrenergic neurons of the locus coeruleus are phase locked to cortical up-down states during sleep. Cereb Cortex 22:426–435.

Fattinger S, De Beukelaar TT, Ruddy KL, Volk C, Heyse NC, Herbst JA, Hahnloser RHR, Wenderoth N, Huber R (2017) Deep sleep maintains learning efficiency of the human brain. Nat Commun 8:1–13.

Fernandez LM, Vantomme G, Osorio-Forero A, Cardis R, Béard E, Lüthi A (2018) Thalamic reticular control of local sleep in mouse sensory cortex. Elife 7.

Fernandez LMJ, Lüthi A (2020) Sleep spindles: Mechanisms and functions. Physiol Rev 100:805–868.

Genzel L, Kroes MCW, Dresler M, Battaglia FP (2014) Light sleep versus slow wave sleep in memory consolidation: A question of global versus local processes? Trends Neurosci 37:10–19.

Hagler DJ, Ulbert I, Wittner L, Erőss L, Madsen JR, Devinsky O, Doyle W, Fabo D, Cash SS, Halgren E (2018) Heterogeneous origins of human sleep spindles in different cortical layers. J Neurosci 38:2241–17.

Halász P (2016) The K-complex as a special reactive sleep slow wave - A theoretical update. Sleep Med Rev 29:34–40.

Hayat H, Regev N, Matosevich N, Sales A, Paredes-Rodriguez E, Krom AJ, Bergman L, Li Y, Lavigne M, Kremer EJ, Yizhar O, Pickering AE, Nir Y (2020) Locus coeruleus norepinephrine activity mediates sensory-evoked awakenings from sleep. Sci Adv 6.

Henin S, Borges H, Shankar A, Sarac C, Melloni L, Friedman D, Flinker A, Parra LC, Buzsaki G, Devinsky O, Liu A (2019) Closed-loop acoustic stimulation enhances sleep oscillations but not memory performance. eneuro ENEURO.0306-19.2019.

Hubbard J, Gent TC, Hoekstra MMB, Emmenegger Y, Mongrain V, Landolt HP, Adamantidis AR, Franken P (2020) Rapid fast-delta decay following prolonged wakefulness marks a phase of wake-inertia in NREM sleep. Nat Commun 11:1–16.

Iber C, Ancoli-Israel S, Chesson Jr. AL, Quan SF (2007) The AASM Manual for the Scoring of Sleep and Associated Events: Rules Terminology and Technical Specifications 1st ed. Westchester, IL: American Academy of Sleep Medicine.

Inostroza M, Born J (2013) Sleep for preserving and transforming episodic memory. Annu Rev Neurosci 36:79–102.

Jiang X, Gonzalez-Martinez J, Halgren E (2019a) Coordination of Human Hippocampal Sharpwave Ripples during NREM Sleep with Cortical Theta Bursts, Spindles, Downstates, and Upstates. J Neurosci 39:8744–8761.

Jiang X, Gonzalez-Martinez J, Halgren E (2019b) Posterior Hippocampal Spindle Ripples Co-occur with Neocortical Theta Bursts and Downstates-Upstates, and Phase-Lock with Parietal Spindles during NREM Sleep in Humans. J Neurosci 39:8949–8968.

Keller-Mcnulty A, Higgins J j (1987) Effect of tail weight and outliers on power and type-i error of robust permutation tests for location. Commun Stat - Simul Comput 16:17–35.

Kim J, Gulati T, Ganguly K (2019) Competing Roles of Slow Oscillations and Delta Waves in Memory Consolidation versus Forgetting. Cell 179:514–526.e13.

Kleiner M, Brainard D, Pelli D, Ingling A, Murray R, Broussard C (2007) What’s new in psychtoolbox-3. Perception 36:1–-16 .

Klinzing JG, Mölle M, Weber FD, Supp G, Hipp JF, Engel AK, Born J (2016) Spindle activity phase-locked to sleep slow oscillations. Neuroimage 134:607–616.

Klinzing JG, Niethard N, Born J (2019) Mechanisms of systems memory consolidation during sleep. Nat Neurosci.

Krugliakova E, Volk C, Jaramillo V, Sousouri G, Huber R (2020) Changes in cross-frequency coupling following closed-loop auditory stimulation in non-rapid eye movement sleep. Sci Rep 10:1–12.

Lacourse K, Delfrate J, Beaudry J, Peppard P, Warby SC (2019) A sleep spindle detection algorithm that emulates human expert spindle scoring. J Neurosci Methods 316:3–11.

Le Van Quyen M, Muller LE, Telenczuk B, Halgren E, Cash S, Hatsopoulos NG, Dehghani N, Destexhe A (2016) High-frequency oscillations in human and monkey neocortex during the wake-sleep cycle. Proc Natl Acad Sci 113:9363–9368.

Le Van Quyen M, Staba R, Bragin A, Dickson C, Valderrama M, Fried I, Engel J (2010) Large-scale microelectrode recordings of high-frequency gamma oscillations in human cortex during sleep. J Neurosci 30:7770–82.

Lecci S, Fernandez LMJ, Weber FD, Cardis R, Chatton JY, Born J, Lüthi A (2017) Coordinated infraslow neural and cardiac oscillations mark fragility and offline periods in mammalian sleep. Sci Adv 3.

Lemieux M, Chen J-Y, Lonjers P, Bazhenov M, Timofeev I (2014) The impact of cortical deafferentation on the neocortical slow oscillation. J Neurosci 34:5689–703.

Leminen M, Virkkala J, Saure E, Paajanen T, Zee P, Santostasi G, Hublin C, Müller K, Porkka-Heiskanen T, Huotilainen M, Paunio T (2017) Enhanced Memory Consolidation Via Automatic Sound Stimulation during Non-REM Sleep. Sleep 40.

Lüthi A (2014) Sleep Spindles: Where They Come From, What They Do. Neurosci 20:243–256.

Maingret N, Girardeau G, Todorova R, Goutierre M, Zugaro M (2016) Hippocampo-cortical coupling mediates memory consolidation during sleep. Nat Neurosci 19:959–964.

Mak-McCully RA, Deiss SR, Rosen BQ, Jung K-Y, Sejnowski TJ, Bastuji H, Rey M, Cash SS, Bazhenov M, Halgren E (2014) Synchronization of isolated downstates (K-complexes) may be caused by cortically-induced disruption of thalamic spindling. PLoS Comput Biol 10:e1003855.

Mak-McCully RA, Rolland M, Sargsyan A, Gonzalez C, Magnin M, Chauvel P, Rey M, Bastuji H, Halgren E (2017) Coordination of cortical and thalamic activity during non-REM sleep in humans. Nat Commun 8:15499.

Maris E, Oostenveld R (2007) Nonparametric statistical testing of EEG-and MEG-data. J Neurosci Methods 164:177–190.

Mensen A, Riedner B, Tononi G (2016) Optimizing detection and analysis of slow waves in sleep EEG. J Neurosci Methods 274:1–12.

Milligan GW, Cooper MC (1988) A study of standardization of variables in cluster analysis. J Classif 5:181–204.

Mukovski M, Chauvette S, Timofeev I, Volgushev M (2007) Detection of active and silent states in neocortical neurons from the field potential signal during slow-wave sleep. Cereb Cortex 17:400–414.

Navarrete M, Arthur S, Treder MS, Lewis PA (2022) Ongoing neural oscillations predict the post-stimulus outcome of closed loop auditory stimulation during slow-wave sleep. Neuroimage 253:119055.

Navarrete M, Schneider J, Ngo H V, Valderrama M, Casson AJ, Lewis PA (2020a) Examining the optimal timing for closed-loop auditory stimulation of slow-wave sleep in young and older adults. Sleep 43:1–14.

Navarrete M, Valderrama M, Lewis PA (2020b) The role of slow-wave sleep rhythms in the cortical-hippocampal loop for memory consolidation. Curr Opin Behav Sci 32:102–110.

Nghiem T-AE, Tort-Colet N, Górski T, Ferrari U, Moghimyfiroozabad S, Goldman JS, Teleńczuk B, Capone C, Bal T, di Volo M, Destexhe A (2020) Cholinergic Switch between Two Types of Slow Waves in Cerebral Cortex. Cereb Cortex 3451–3466.

Ngo H-V V, Martinetz T, Born J, Mölle M (2013) Auditory closed-loop stimulation of the sleep slow oscillation enhances memory. Neuron 78:545–53.

Ngo H-V V, Miedema A, Faude I, Martinetz T, Molle M, Born J (2015) Driving Sleep Slow Oscillations by Auditory Closed-Loop Stimulation--A Self-Limiting Process. J Neurosci 35:6630–6638.

Nichols TE, Holmes AP (2002) Nonparametric permutation tests for functional neuroimaging: A primer with examples. Hum Brain Mapp 15:1–25.

Nir Y, Staba RJ, Andrillon T, Vyazovskiy V V, Cirelli C, Fried I, Tononi G (2011) Regional Slow Waves and Spindles in Human Sleep. Neuron 70:153–169.

Ong JL, Lo JC, Chee NIYN, Santostasi G, Paller KA, Zee PC, Chee MWLL (2016) Effects of phase-locked acoustic stimulation during a nap on EEG spectra and declarative memory consolidation. Sleep Med 20:88–97.

Osorio-Forero A, Cardis R, Vantomme G, Guillaume-Gentil A, Katsioudi G, Devenoges C, Fernandez LMJ, Lüthi A (2021) Noradrenergic circuit control of non-REM sleep substates. Curr Biol 31:5009–5023.e7.

Papalambros NA, Santostasi G, Malkani RG, Braun R, Weintraub S, Paller KA, Zee PC (2017) Acoustic enhancement of sleep slow oscillations and concomitant memory improvement in older adults. Front Hum Neurosci 11:1–14.

Pelli DG (1997) The VideoToolbox software for visual psychophysics: transforming numbers into movies. Spat Vis.

Purcell SM, Manoach DS, Demanuele C, Cade BE, Mariani S, Cox R, Panagiotaropoulou G, Saxena R, Pan JQ, Smoller JW, Redline S, Stickgold R (2017) Characterizing sleep spindles in 11,630 individuals from the National Sleep Research Resource. Nat Commun 8:15930.

Rasch B, Born J (2013) About sleep’s role in memory. Physiol Rev 93:681–766.

Riedner B a, Vyazovskiy V V, Huber R, Massimini M, Esser S, Murphy M, Tononi G (2007) Sleep Homeostasis and Cortical Synchronization: III. A High-Density EEG Study of Sleep Slow Waves in Humans. Sleep 30:1643–1657.

Rosanova M, Timofeev I (2005) Neuronal mechanisms mediating the variability of somatosensory evoked potentials during sleep oscillations in cats. J Physiol 562:569–582.

Sauseng P, Klimesch W, Gruber WR, Hanslmayr S, Freunberger R, Doppelmayr M (2007) Are event-related potential components generated by phase resetting of brain oscillations? A critical discussion. Neuroscience 146:1435–1444.

Schabus M, Dang-Vu TT, Heib DPJ, Boly M, Desseilles M, Vandewalle G, Schmidt C, Albouy G, Darsaud A, Gais S, Degueldre C, Balteau E, Phillips C, Luxen A, Maquet P (2012) The Fate of Incoming Stimuli during NREM Sleep is Determined by Spindles and the Phase of the Slow Oscillation. Front Neurol 3:40.

Schneider J, Lewis PA, Koester D, Born J, Ngo H-V V (2020) Susceptibility to auditory closed-loop stimulation of sleep slow oscillations changes with age. Sleep 1–10.

Siclari F, Bernardi G, Riedner B a, LaRocque JJ, Benca RM, Tononi G (2014) Two Distinct Synchronization Processes in the Transition to Sleep: A High-Density Electroencephalographic Study. Sleep 37:1621–1637.

Staresina BP, Ole Bergmann T, Bonnefond M, van der Meij R, Jensen O, Deuker L, Elger CE, Axmacher N, Fell J (2015) Hierarchical nesting of slow oscillations, spindles and ripples in the human hippocampus during sleep. Nat Neurosci 18:1679–1686.

Steriade M (2003) The corticothalamic system in sleep. Front Biosci 8:d878–99.

Steriade M, Contreras D, Curró Dossi R, Nuñez A (1993a) The slow (<1 Hz) oscillation in reticular thalamic and thalamocortical neurons: scenario of sleep rhythm generation in interacting thalamic and neocortical networks. J Neurosci 13:3284–99.

Steriade M, McCormick DA, Sejnowski T (1993b) Thalamocortical oscillations in the sleeping and aroused brain. Science (80-) 262:679–685.

Steriade M, Nunez a, Amzica F (1993c) Intracellular analysis of relations between the slow (< 1 Hz) neocortical oscillation and other sleep rhythms of the electroencephalogram. J Neurosci 13:3266–3283.

Todorova R, Zugaro M (2020) Hippocampal ripples as a mode of communication with cortical and subcortical areas. Hippocampus 30:39–49.

Torres FA, Orio P, Escobar MJ (2021) Selection of stimulus parameters for enhancing slow wave sleep events with a neural-field theory thalamocortical model. PLoS Comput Biol 17:1–28.

Vyazovskiy V V., Olcese U, Lazimy YM, Faraguna U, Esser SK, Williams JC, Cirelli C, Tononi G (2009) Cortical Firing and Sleep Homeostasis. Neuron 63:865–878.

Warby SC, Wendt SL, Welinder P, Munk EGS, Carrillo O, Sorensen HBD, Jennum P, Peppard PE, Perona P, Mignot E (2014) Sleep-spindle detection: crowdsourcing and evaluating performance of experts, non-experts and automated methods. Nat Methods 11:385–392.

Wei Y, Krishnan GP, Marshall L, Martinetz T, Bazhenov M (2020) Stimulation Augments Spike Sequence Replay and Memory Consolidation during Slow-Wave Sleep. J Neurosci 40:811–824.

