## Supplemental Files for "Response of sleep slow oscillations to acoustic stimulation is evidenced by distinctive synchronization processes"

### Supplementary Figures

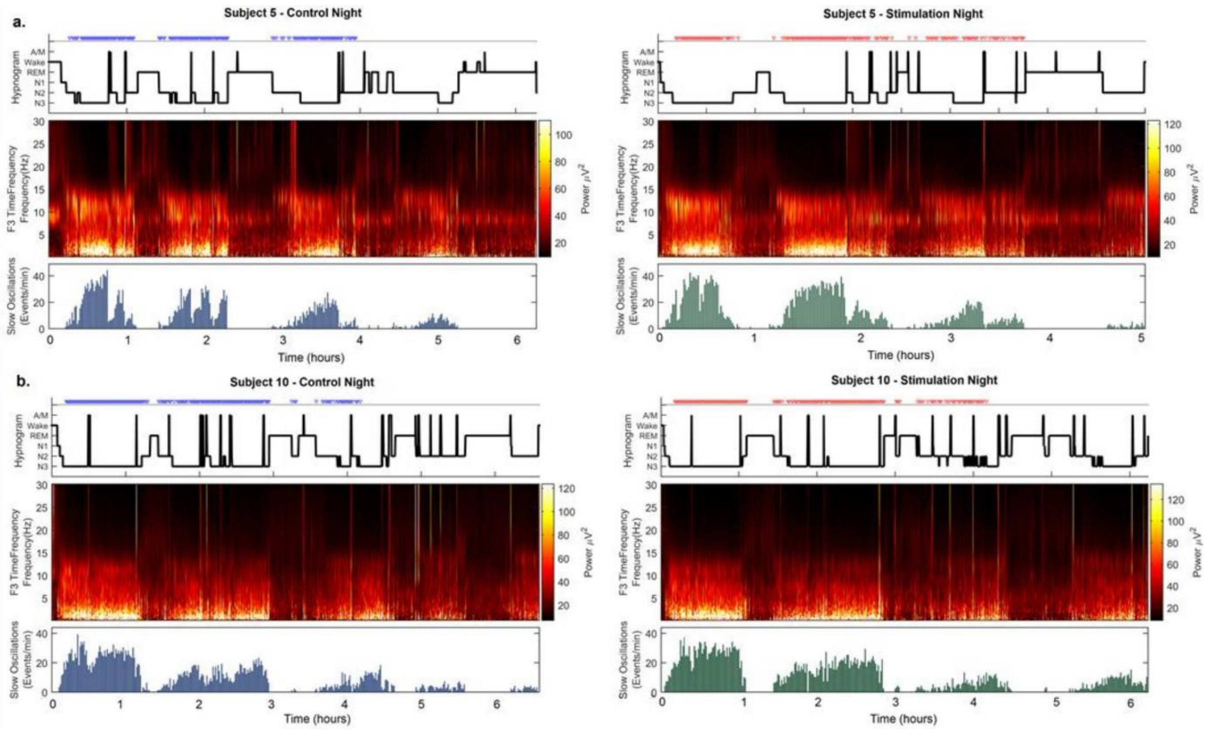


Figure S1. Hypnograms for two representative subjects. (a) and (b) present the hypnogram for control (left) and stimulation (right) nights. Time-frequency plots demonstrate oscillatory activity according to sleep stages and SWS activity from the detected SO density. Marked clicks are visualized above the hypnograms (solid blue or red bars) but were not shown during scoring.

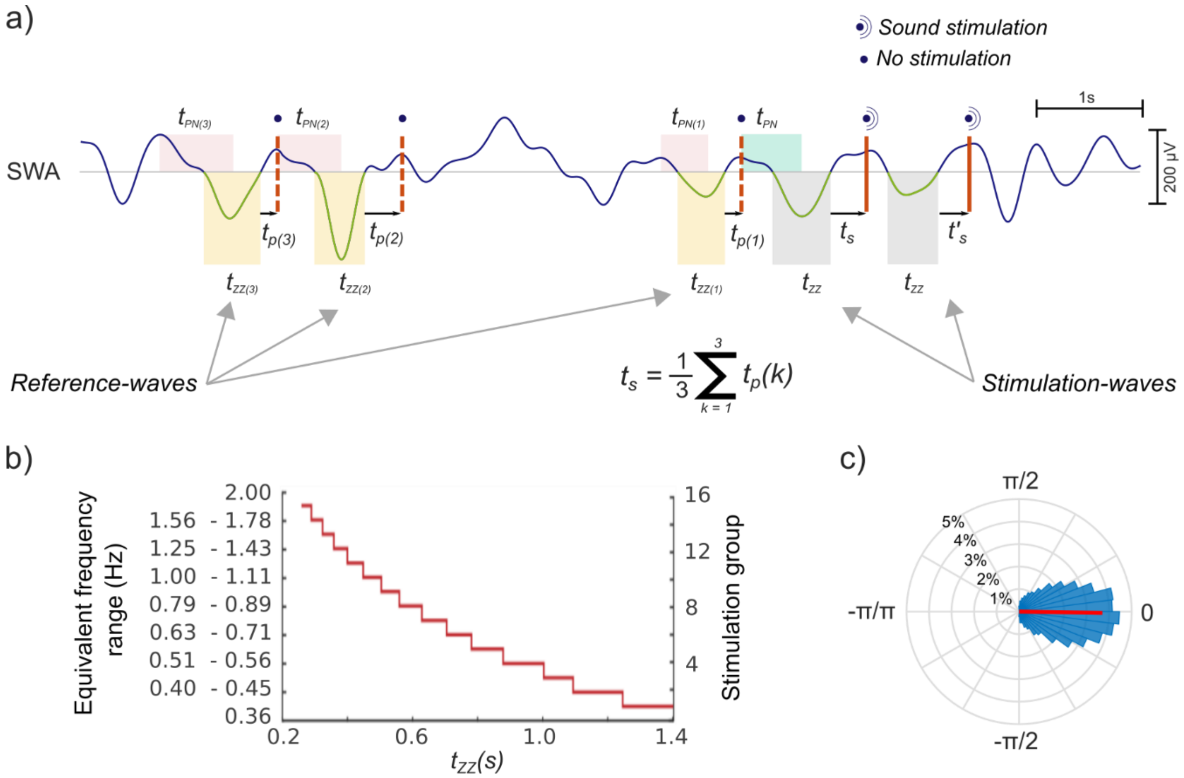


Figure S2. Online detection and stimulation of SO. (a) online SO morphology detection. Only waves with $t_{PN}$ and $t_{ZZ}$ times in the SO interval (~250 to 1400 ms) were considered SO. For pre-defined frequency intervals, the last three $t_P$ times detected were saved into memory from detection of non-stimulated reference waves. The time of stimulation was computed as the average $t_P$ time to be used as $t_S$ on stimulation waves. Two clicks were applied following the same method and then the stimulation was paused for 2.5 seconds. SO detections during this period of non-stimulation were thus used as reference waves (b) Detected waves were classified in fifteen frequency intervals (stimulation groups) depending on the negative deflection period ($t_{ZZ}$). (c) Click distributions and average phase of stimulation for all stimulated SO in CNT and STM conditions.

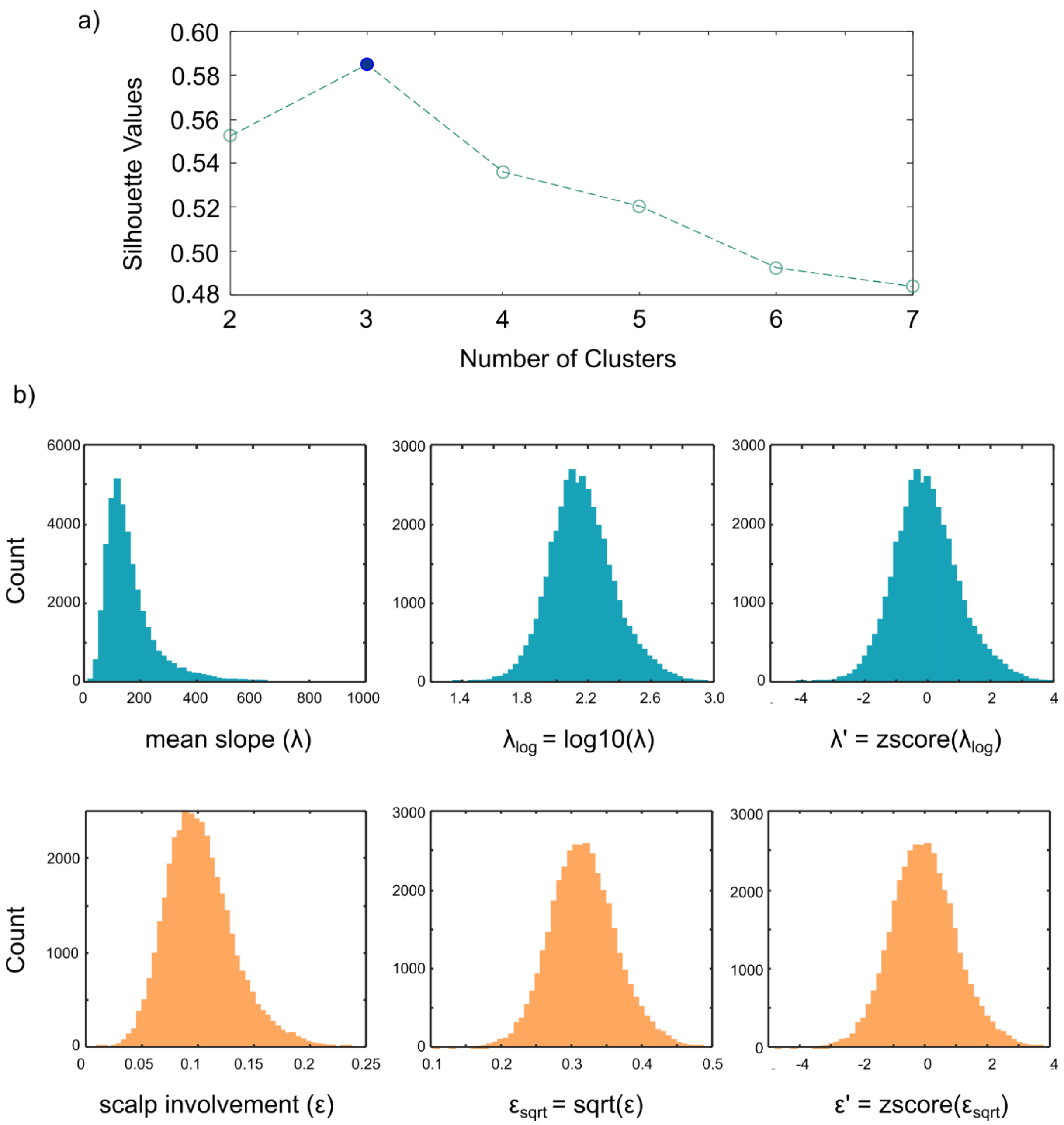


Figure S3. Pre-processing for k-means clustering analysis. (a) A silhouette analysis to study the separation distance between the resulting clusters indicated that the optimal grouping is three clusters. (b) Distributions for mean slope (λ) and scalp involvement (ε) indicated non-normality for both parameters. Therefore, both parameters were normalized and then z scored. This pre-processing procedure was implemented to prevent any distortion of the relative nearness of observations by the units of each dimension.

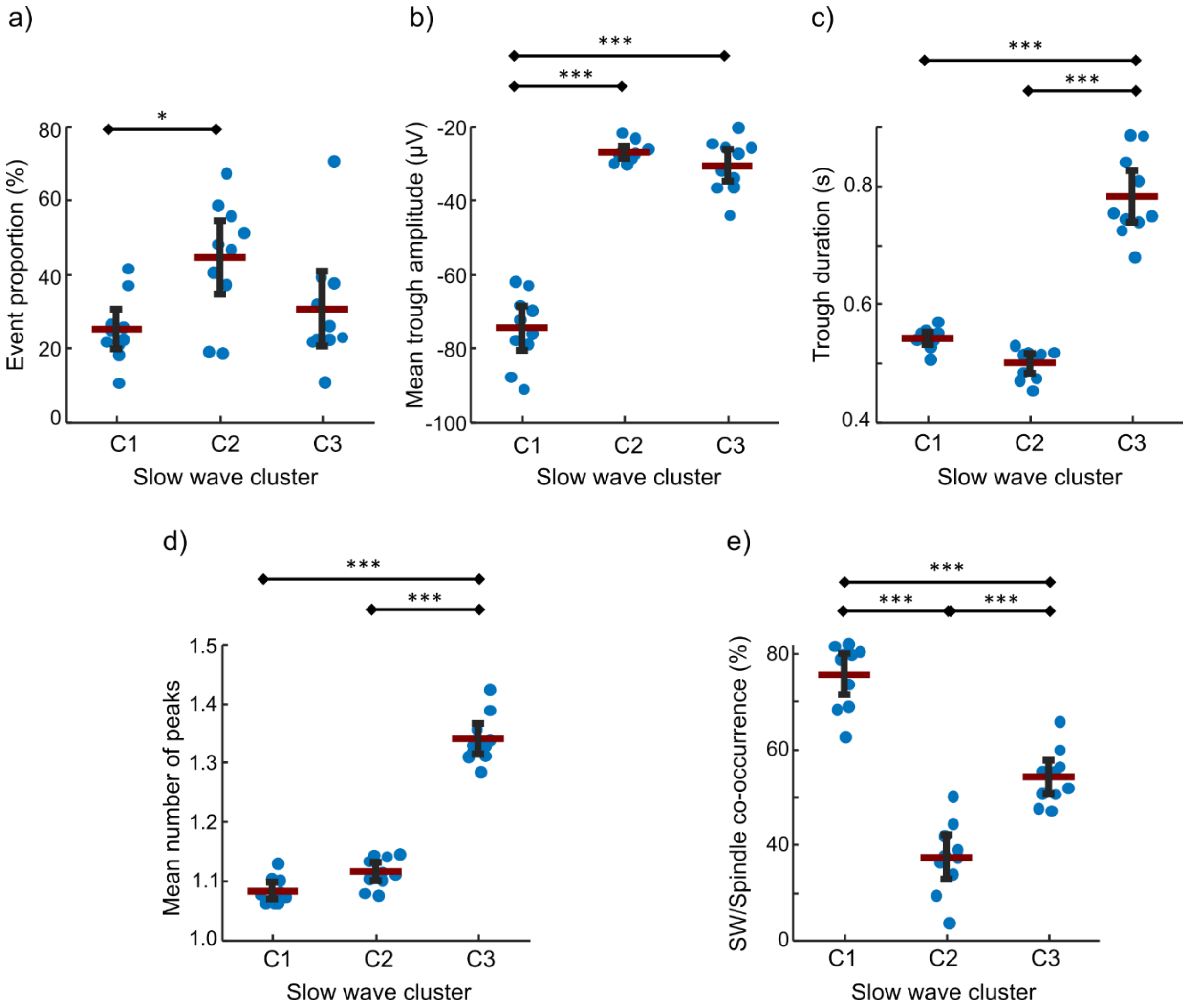


Figure S4. Differences in characteristics between spontaneous slow wave categories averaged across electrodes during N2. (a) Event proportion. (b) Trough amplitude. (c) Trough duration. (d) Number of negative peaks. (e) Event co-occurrence with detected sleep spindles.

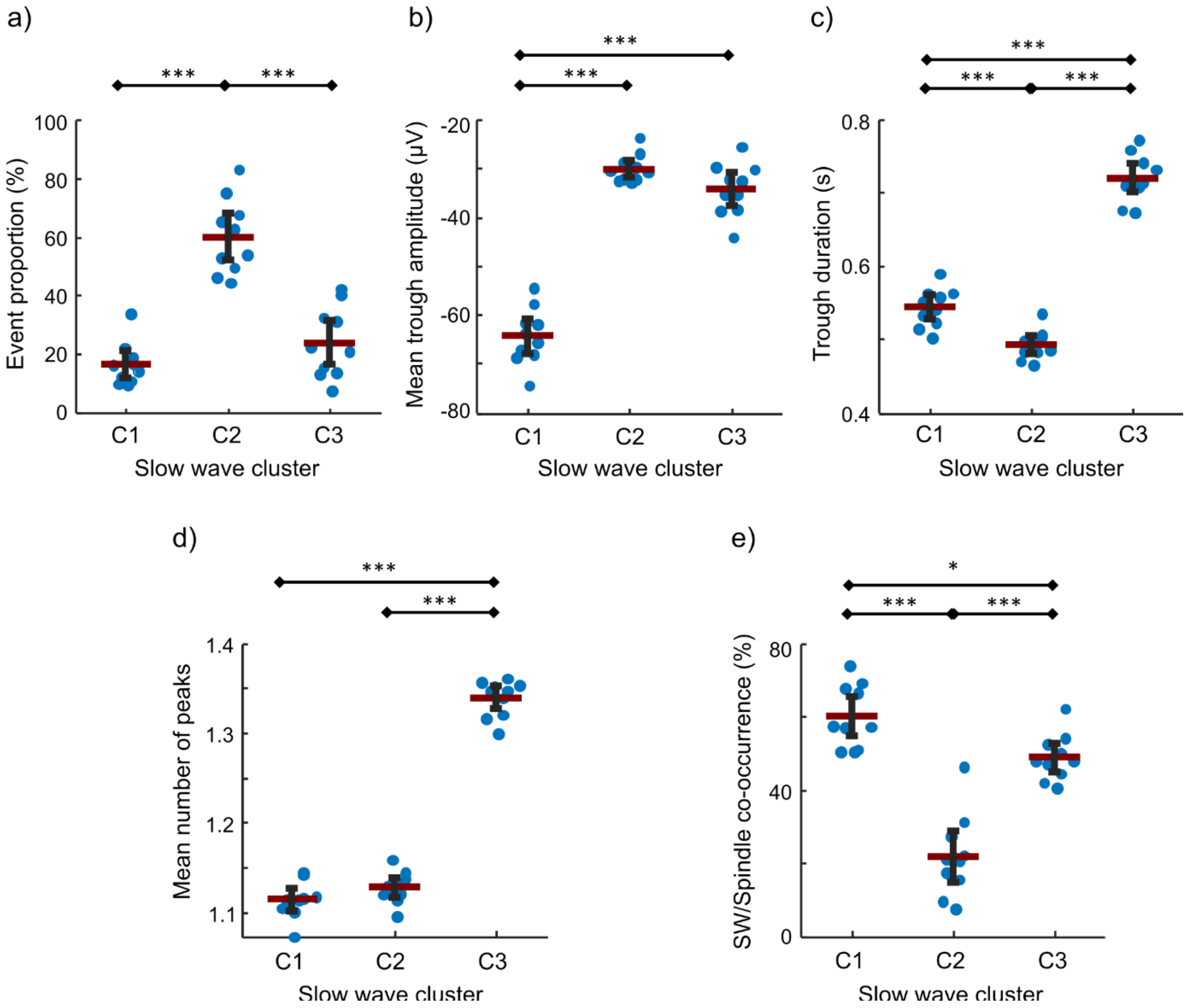


Figure S5. Differences in characteristics between spontaneous slow wave categories averaged across electrodes during N3. (a) Event proportion. (b) Trough amplitude. (c) Trough duration. (d) Number of negative peaks. (e) Event co-occurrence with detected sleep spindles.

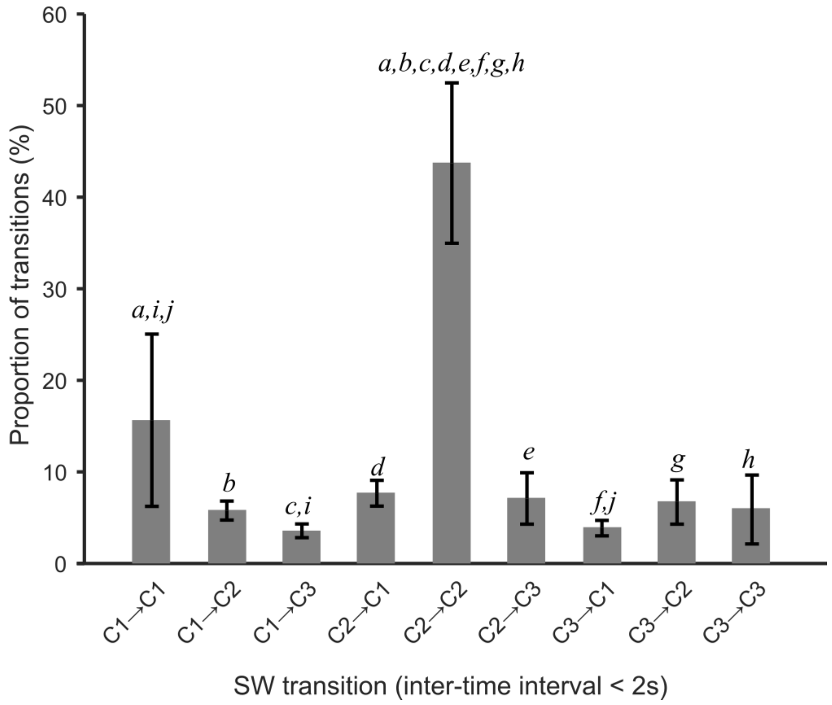


Figure S6. Proportion of event transitions between the different SW categories averaged across participants. SW transitions were represented as (first event → following event) for SW spaced out less than 2 seconds. Small italic letters represent pairs of transitions with significant differences in proportion after post-hoc analysis.

**Supplementary Tables**

**Table S1.** Sleep architecture

| Variable | Control (mean ± SD) | Stimulation (mean ± SD) | *t* | *p* |
|---|---|---|---|---|
| *Recording time* | 407.07 ± 36.21 | 400.54 ± 49.35 | -0.87 | 0.402 |
| *Efficiency (%)* | 95.73 ± 5.25 | 96.38 ± 3.94 | 0.50 | 0.628 |
| *TST* | 389.25 ± 36.85 | 385.43 ± 45.59 | -0.45 | 0.657 |
| *Sleep period* | 399.11 ± 33.74 | 392.39 ± 46.35 | -0.92 | 0.373 |
| *Awakenings (#)* | 3.86 ± 1.99 | 4.93 ± 3.95 | 1.72 | 0.110 |
| *Sleep latency* | 7.96 ± 8.72 | 8.14 ± 14.65 | 0.05 | 0.960 |
| *WASO Time* | 9.86 ± 16.15 | 6.96 ± 8.79 | -0.91 | 0.380 |
| *N1 time* | 15.14 ± 13.51 | 19.32 ± 18.72 | 1.39 | 0.189 |
| *N2 time* | 152.36 ± 29.73 | 147.18 ± 27.77 | -0.77 | 0.455 |
| *N3 time* | 96.61 ± 46.84 | 102.07 ± 47.20 | 1.43 | 0.177 |
| *REM time* | 81.89 ± 22.50 | 73.43 ± 29.74 | -1.18 | 0.258 |

**Note:** All times are expressed in minutes. TST: Total sleep time. WASO: wake after sleep onset. SD: standard deviation

**Table S2.** Morphological features of SW categories across all the night and between conditions

| Variable | CNT | | STM | | *t* | *p* |
|---|---|---|---|---|---|---|
| | mean | SD | mean | SD | | |
| ***C1 events*** | | | | | | |
| *SW proportion (%)* | 25.78 | 19.75 | 28.30 | 16.67 | -0.85 | 0.409 |
| *Trough amplitude (μV)* | -70.20 | 5.90 | -73.86 | 12.43 | 1.44 | 0.174 |
| *Trough duration (s)* | 0.54 | 0.02 | 0.54 | 0.02 | -0.63 | 0.542 |
| *Number of peaks* | 1.10 | 0.02 | 1.10 | 0.02 | -0.49 | 0.630 |
| *SW/Spindle co-occurrence (%)* | 65.70 | 8.66 | 64.17 | 9.11 | 0.99 | 0.338 |
| *SW density (z-score)* | | | | | | |
| *1st NREM cycle* | -0.83 | 0.37 | -0.87 | 0.31 | 0.55 | 0.590 |
| *2nd NREM cycle* | -0.81 | 0.57 | -0.89 | 0.31 | 0.52 | 0.611 |
| *3rd NREM cycle* | -0.88 | 0.33 | -0.86 | 0.29 | -0.20 | 0.844 |
| *4th NREM cycle* | -0.93 | 0.28 | -0.88 | 0.26 | -1.09 | 0.303 |
| ***C2 events*** | | | | | | |
| *SW proportion (%)* | 53.00 | 16.33 | 51.79 | 14.30 | 0.37 | 0.718 |
| *Trough amplitude (μV)* | -31.60 | 4.92 | -33.06 | 5.38 | 2.08 | 0.058 |
| *Trough duration (s)* | 0.50 | 0.02 | 0.50 | 0.03 | -0.76 | 0.460 |
| *Number of peaks* | 1.12 | 0.02 | 1.12 | 0.03 | -0.37 | 0.717 |
| *SW/Spindle co-occurrence (%)* | 22.10 | 10.97 | 18.01 | 13.21 | 1.82 | 0.091 |
| *SW density (z-score)* | | | | | | |
| *1st NREM cycle* | -0.22 | 0.49 | -0.35 | 0.45 | 0.80 | 0.437 |
| *2nd NREM cycle* | -0.56 | 0.38 | -0.53 | 0.36 | -0.18 | 0.863 |
| *3rd NREM cycle* | -0.68 | 0.36 | -0.72 | 0.32 | 0.33 | 0.747 |
| *4th NREM cycle* | -0.91 | 0.20 | -0.93 | 0.17 | -0.35 | 0.737 |
| ***C3 events*** | | | | | | |
| *SW proportion (%)* | 21.22 | 14.22 | 19.91 | 14.56 | 0.57 | 0.581 |
| *Trough amplitude (μV)* | -35.41 | 6.91 | -39.94 | 10.72 | 2.02 | 0.064 |
| *Trough duration (s)* | 0.74 | 0.04 | 0.76 | 0.07 | -1.23 | 0.241 |
| *Number of peaks* | 1.33 | 0.03 | 1.34 | 0.03 | -1.37 | 0.195 |
| *SW/Spindle co-occurrence (%)* | 52.74 | 7.85 | 56.23 | 14.90 | -0.85 | 0.411 |
| *SW density (z-score)* | | | | | | |
| *1st NREM cycle* | -0.91 | 0.19 | -0.96 | 0.25 | 0.97 | 0.348 |
| *2nd NREM cycle* | -0.98 | 0.20 | -1.00 | 0.21 | 0.21 | 0.836 |
| *3rd NREM cycle* | -0.93 | 0.22 | -1.02 | 0.11 | 1.57 | 0.140 |
| *4th NREM cycle* | -1.01 | 0.25 | -0.94 | 0.21 | -1.92 | 0.087 |
